# Orthogonal coupling of a 3D cytoskeletal scaffold coordinates cell morphogenesis and maintains tissue organization in the Drosophila pupal retina

**DOI:** 10.1101/2023.03.06.531386

**Authors:** Xiao Sun, Jacob Decker, Nicelio Sanchez-Luege, Ilaria Rebay

**Affiliations:** Committee on Development, Regeneration and Stem Cell Biology; Ben May Department for Cancer Research University of Chicago, Chicago, IL 60637 USA; Corresponding Author

**Keywords:** Abelson, Drosophila eye development, morphogenesis, 3D cytoskeletal network, apical-basal polarity, actin cytoskeleton, epithelial patterning, cell-cell interactions, feedback mechanism, photoreceptor, pigment cell

## Abstract

How complex three-dimensional (3D) organs coordinate cellular morphogenetic events to achieve the correct final form is a central question in development. The question is uniquely tractable in the late *Drosophila* pupal retina where cells maintain stereotyped contacts as they elaborate the specialized cytoskeletal structures that pattern the apical, basal and longitudinal planes of the epithelium. In this study, we combined cell type-specific genetic manipulation of the cytoskeletal regulator Abelson (Abl) with 3D imaging to explore how the distinct cellular morphogenetic programs of photoreceptors and interommatidial pigment cells coordinately organize tissue pattern to support retinal integrity. Our experiments revealed an unanticipated intercellular feedback mechanism whereby correct cellular differentiation of either cell type can non-autonomously induce cytoskeletal remodeling in the other *Abl* mutant cell type, restoring retinal pattern and integrity. We propose that genetic regulation of specialized cellular differentiation programs combined with inter-plane mechanical feedback confers spatial coordination to achieve robust 3D tissue morphogenesis.

## Introduction

The spatial arrangement of cells within an epithelium is critical to the final form and function of the tissue. During development, genetically controlled terminal differentiation programs produce the specialized cytoskeletal structures, cell-cell junctional adhesions and cell-extracellular matrix (ECM) contacts unique to each cell type. In turn, the resulting cell shapes, structures and connections introduce specific packing constraints that influence final organ form. While progress has been made in describing the acquisition of tissue form in simple epithelia with relatively homogeneous cell composition, how complex tissues with diverse cell fates, shapes, and physical properties spatially coordinate dramatic morphogenetic remodeling to maintain robust organization remains poorly understood (Collinet and Lecuit, 2021).

Previous studies have shown that coordinated cell shape changes driven by subcellular cytoskeletal and junction remodeling produce tissue-level morphology. The best-studied process is apical constriction, where supracellular networks physically couple cell apices across a tissue plane to drive numerous morphogenetic processes including epithelial folding, bending, invagination and closure (Martin and Goldstein, 2014; Perez-Vale and Peifer, 2020). Subsequent consideration of cells as 3D units has emphasized how remodeling of cellular structure along basal or lateral planes or of the ECM can also promote morphogenetic change (Daley and Yamada, 2013; Gracia et al., 2019; Harmansa et al., 2022; Roellig et al., 2022; Sui et al., 2018). Temporal sequences of independent planar changes also coordinate 3D change. Notable examples include the sequential activation of actomyosin contractility along different planes to organize the successive patterns of apical and lateral contraction required for endoderm invagination in the ascidian embryo (Sherrard et al., 2010) and lumen morphogenesis in the C. elegans vulva (Yang et al., 2017). In all these examples, the apical-lateral-basal organization inherent to polarized epithelial tissues provides an intuitive physical conduit for driving 3D cellular and tissue-level morphogenetic change. Despite the established importance of supracellular networks in providing mechanical coupling within individual planes, whether and how remodeling processes interact across different planes to produce specific 3D cellular and tissue scale morphologies remains to be explored.

The stereotyped pseudostratified epithelial architecture of the Drosophila compound eye makes it an attractive model to approach this question. The fly retina is a complex epithelial organ whose form and function depends on the precise organization of highly specialized and uniquely shaped cell types (Figure 1A and 1B; (Charlton-Perkins and Cook, 2010; Ready et al., 1976a; Wolff and Ready, 1993). Clusters of eight photoreceptor neurons occupy the central core of each ommatidial unit, with their photosensitive rhabdomeres defining the longitudinal optical axis of the epithelium. Directly above each cluster, an apical assembly of four cone and two primary pigment cells produces the lens that will focus incoming light onto the underlying photoreceptors. A hexagonal lattice of secondary and tertiary interommatidial pigment cells (IOPCs) surrounds each photoreceptor cluster. Through their cortical cytoskeletal-enriched junctional domains and basal cell-cell/ECM junctional contacts, the IOPCs provide in-plane connections across the apical and basal planes of the retinal field.

**Figure 1.**
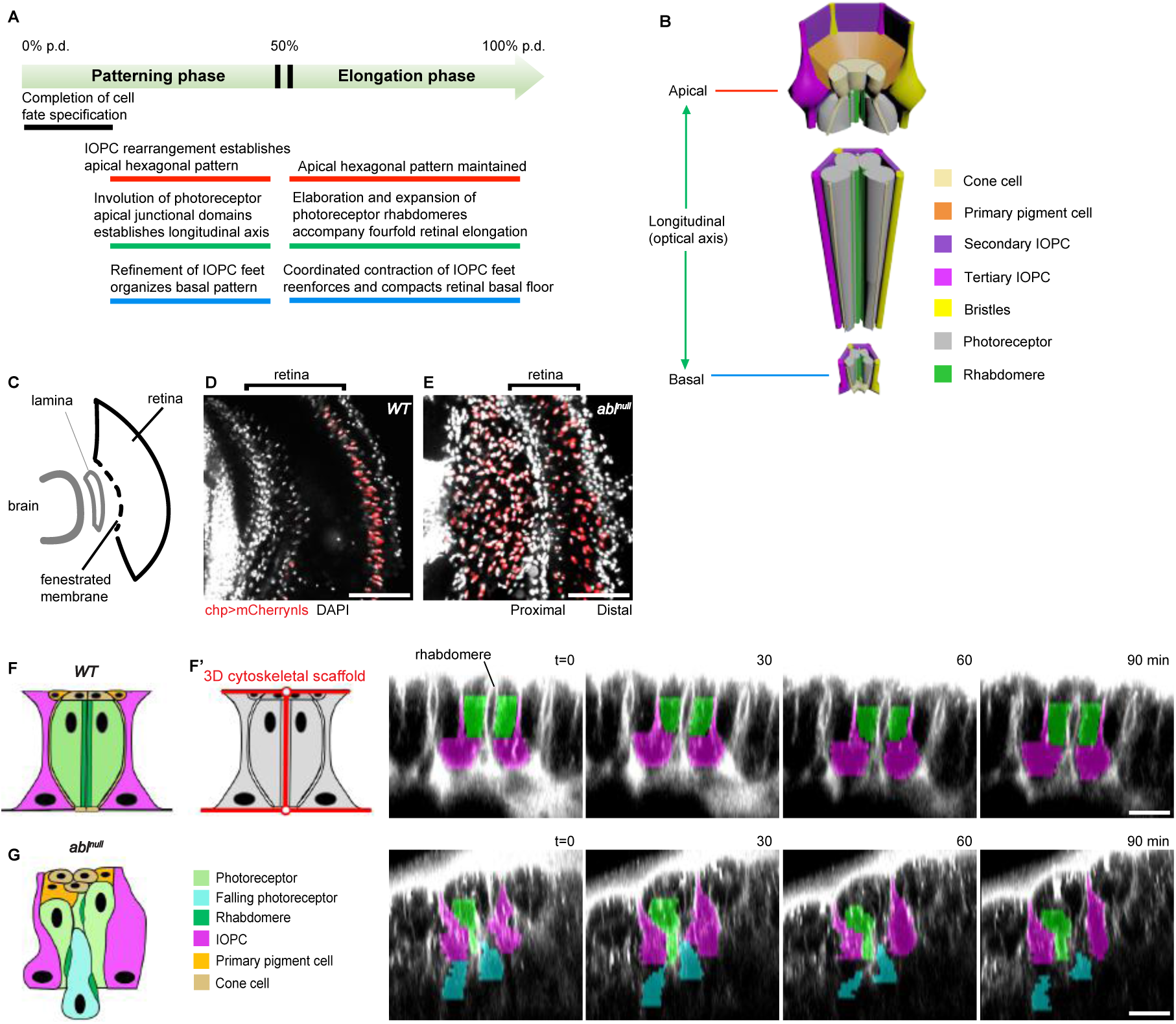
Loss of *abl* results in photoreceptors “falling” out of the retinal epithelium. **(A)** Timeline summarizing the sequence of key morphogenetic events that pattern the apical, longitudinal and basal planes of the pupal retinal epithelium. IOPC, interommatidial pigment cell; p.d., pupal development. **(B)** Schematic summarizing the 3D organization of a 100% p.d. ommatidium. **(C)** Schematic of the adult visual system. The retinal fenestrated membrane (dashed line) separates it from the underlying lamina, the distal-most ganglion of the brain optic lobe. **(D,E)** Comparison of photoreceptor nuclear position (red) in 100% p.d. retinal-brain complexes. DAPI (white) marks all nuclei. Scale bar = 50 µm. **(F, G.)** Schematics and stills from time-lapse movies (Videos S1 and S2) of 50% p.d retinas injected with CellMask (white). False color shows photoreceptors (green), fallen photoreceptors (cyan) and secondary IOPCs (magenta) in a representative ommatidium. Scale bars = 10µm. See also Figure S2A-S2D. **F’** depicts the 3D scaffold.

Pupal retinal development can be separated into two phases, a pattern establishment phase and a tissue elongation phase during which pattern is maintained (Figure 1A)(Cagan and Ready, 1989; Johnson, 2021; Ready et al., 1976b). During the patterning phase, cone cell and IOPC rearrangements produce the precise hexagonal lattice pattern while photoreceptor apical domain involution and anchorage to the cone cells aligns the optical axis relative to the apical and basal surfaces. During the elongation phase, cell-cell contacts and overall tissue organization are maintained while the photoreceptor and IOPC terminal differentiation programs elaborate functional specialized structures and the epithelium elongates four-fold. Although the emergence of two-dimensional (2D) planar pattern and the individual morphogenetic events in different retinal cell types during the early patterning phase have been well-described (Bao and Cagan, 2005; Baumann, 2004; Cagan and Ready, 1989; Hayashi and Carthew, 2004; Hilgenfeldt et al., 2008; Johnson, 2021; Kafer et al., 2007; Longley and Ready, 1995; Pellikka et al., 2002; Pham et al., 2008; Ready, 2002; Ready and Chang, 2021; Signore et al., 2018), how the subsequent cellular morphogenetic changes are regulated, coordinated and integrated across different tissue planes to maintain 3D retinal organization and integrity during retinal elongation has not been explored.

We chose the cytoplasmic tyrosine kinase and cytoskeletal regulator Abelson (Abl) as a tool to examine the impact of modulating retinal cell shapes and structures on tissue organization. Prior phenotypic analyses showed that Abl is required for multiple aspects of the photoreceptor terminal differentiation program and that its loss perturbs ommatidial organization and retinal pattern (Bennett and Hoffmann, 1992; Henkemeyer et al., 1987; Henkemeyer et al., 1990; Kannan et al., 2014; Singh et al., 2010; Xiong and Rebay, 2011; Xiong et al., 2013). Mechanistic studies have shown how Abl modulates cytoskeletal remodeling and junctional dynamics that control cell morphology in a wide range of epithelial tissues. For example, in the early embryonic epithelium Abl helps to coordinate apical constriction during mesoderm invagination and to produce the cell shape changes that drive convergent extension during germband elongation (Fox and Peifer, 2007; Jodoin and Martin, 2016; Tamada et al., 2012; Yu and Zallen, 2020). Inhibition of Enabled (Ena)-mediated linear F-actin assembly is a key aspect of Abl function in many contexts (Comer et al., 1998; Forsthoefel et al., 2005; Fox and Peifer, 2007; Gates et al., 2007; Gertler et al., 1995; Grevengoed et al., 2001; Kannan et al., 2014; Kannan et al., 2017; Lin et al., 2009; Rogers et al., 2021). Best studied is the embryonic central nervous system, where in response to different axon guidance cues, Abl modulates the balance between linear and branched F-actin by regulating the activity not only of Enabled, but also of the WAVE/SCAR complex (Forsthoefel et al., 2005; Gertler et al., 1995; Kannan et al., 2017; Liebl et al., 2000; Wills et al., 1999).

In this study, we investigated how cellular morphogenetic events are individually controlled and effectively communicated between two different retinal cell types as the late pupal retina elaborates and maintains its precise 3D organization. Our approach was to combine genetic perturbation of Abl function with single-cell resolution fixed and live time-lapse imaging to examine retinal cell shapes and tissue-scale patterns. First, we used global depletion of Abl to characterize its contributions to the cytoskeletal specializations each cell type elaborates. In contrast to a wildtype retina where correctly elaborated and aligned photoreceptor and IOPC cytoskeletal domains confine cell shape and organization, loss of Abl disrupted photoreceptor and IOPC terminal differentiation, resulting in a tissue whose heterogenous cellular shapes, structures and intercellular connections were insufficient to maintain retinal integrity. Second, we probed how IOPCs and photoreceptors interact by restoring Abl to each individual cell type in an otherwise *abl* mutant background. Strikingly, these experiments uncovered a feedback interaction between the two cell types that enabled rescue of either photoreceptor or IOPC cellular differentiation to induce morphogenetic remodeling of the other cell type and thus restore retinal 3D organization and tissue integrity. Together our results suggest that the cytoskeletal and junctional adhesions of the photoreceptors and IOPCs provide a mechanically coupled structural scaffold that coordinates cellular differentiation to ensure the robust tissue-level architecture needed for vision.

## Results

### Loss of *abl* results in photoreceptors “falling” out of the retinal epithelium

Prior work showed that in *abl* mutant retinas, photoreceptors are specified correctly but then fail to carry out their normal terminal differentiation program (Henkemeyer et al., 1987; Henkemeyer et al., 1990; Singh et al., 2010; Xiong and Rebay, 2011; Xiong et al., 2013). We were interested in how retinal cell terminal differentiation impacts the elaboration of cell and tissue morphology. As a framework for exploring this, we briefly summarize how the apical, basal and longitudinal networks of actin-based cytoskeletal structures and junctional adhesions established during the patterning phase are subsequently remodeled to produce the specialized structures of a mature wildtype retina (Longley and Ready, 1995; Ready, 2002).

First, the apical network is defined by the hexagonally arranged IOPC apical junctional domains. After its establishment by 50% p.d., the apical network is stably maintained with minimal change (Bao and Cagan, 2005; Cagan and Ready, 1989; Hayashi and Carthew, 2004; Signore et al., 2018). Second, at the basal epithelial plane, IOPCs refine their cell-ECM contacts by anchoring to integrin-reinforced rings, or grommets, at the center of each ommatidium, and by tiling the entire retinal floor with a radial pattern of contractile actomyosin “feet”; during the elongation phase, coordinated contraction of this basal network compacts the retinal floor (Baumann, 2004; Longley and Ready, 1995; Ready and Chang, 2021). Referred to as the “fenestrated membrane”, this specialized contractile network supports and separates the retina from the brain. Third, the longitudinal network is defined by rhabdomeric precursors consisting of the photoreceptor involuted apical membranes which bridge the apical and basal tissue planes at the centerpoint of each ommatidium through junctional connections with the cone cell apical caps and basal feet; for simplicity, we refer to these specialized domains as rhabdomeres, regardless of stage. This unique coupling is established by 50% p.d. and then maintained as the rhabdomeres mature and expand during the elongation phase (Cagan and Ready, 1989). Because rhabdomeres anchor to the cone cell feet, which like the IOPC feet are anchored to the grommets, these rings mark a hub of junctional attachments that physically connect the longitudinal and basal networks and that couple the basal network to the ECM (Figure 1B and 1F). Photoreceptors do not contribute directly to the 2D apical and basal patterns because their longitudinal anchor points are located below and above the apical network plane and basal network plane, respectively. Conversely, IOPCs contribute minimally to the longitudinal network because their thin cellular projections occupy minimal tissue space and do not make junctional connections with photoreceptors along the longitudinal axis.

Using this framework, we first examined final retinal organization by comparing photoreceptor position in wildtype and *abl^null^* 100% p.d. retina-brain complexes (Figure 1C-1E; Figure S1A and S1B). In wildtype, the photoreceptor nuclei clustered in a tight row just below the retinal surface (Figure 1D; Figure S1C and S1D). In contrast, in *abl^null^*, photoreceptor nuclei were scattered throughout the longitudinal length of the retina and were also found fallen beneath the retinal floor in the space between retina and lamina, a phenotype we refer to as loss of retinal integrity. (Figure 1E). Despite the aberrant position of these cells, lineage tracing confirmed their photoreceptor origin and identity (Figure S1E and S1F). Retinal depth was noticeably reduced relative to wildtype (Figure 1E vs. 1D). Thus, Abl is required for photoreceptors to maintain their proper position and for the retina to maintain structural integrity and elongate.

To examine the changes in retinal cell shapes and relative positions during the falling process, we labeled cell membranes and performed time-lapse live imaging (Figure 1F and 1G; Figure S2A-S2D; Videos S1 and S2). In wildtype, completion of the early patterning phase resulted in stereotyped cell shapes, positions and contacts that did not appreciably change (Figure 1F; Figure S2A and S2B). Although the intensity of the injected dye precluded examination of apical network relationships, lateral views highlighted the distinctive interlocking shapes of the photoreceptors and IOPCs (Figure 1F). Photoreceptor cell bodies surround the central rhabdomere-defined longitudinal axis of each ommatidium, occupying the bulk of the tissue space except toward the basal plane, where they narrow to accommodate the IOPC bodies and basal feet. IOPCs maintain connection with the apical plane via thin processes that separate neighboring photoreceptor clusters and fill up the inter-ommatidial space. Despite the lack of junctional connections between photoreceptors and IOPCs, each appears to support and constrain the other’s shape and position through their 3D spatial relationships.

**Figure 2.**
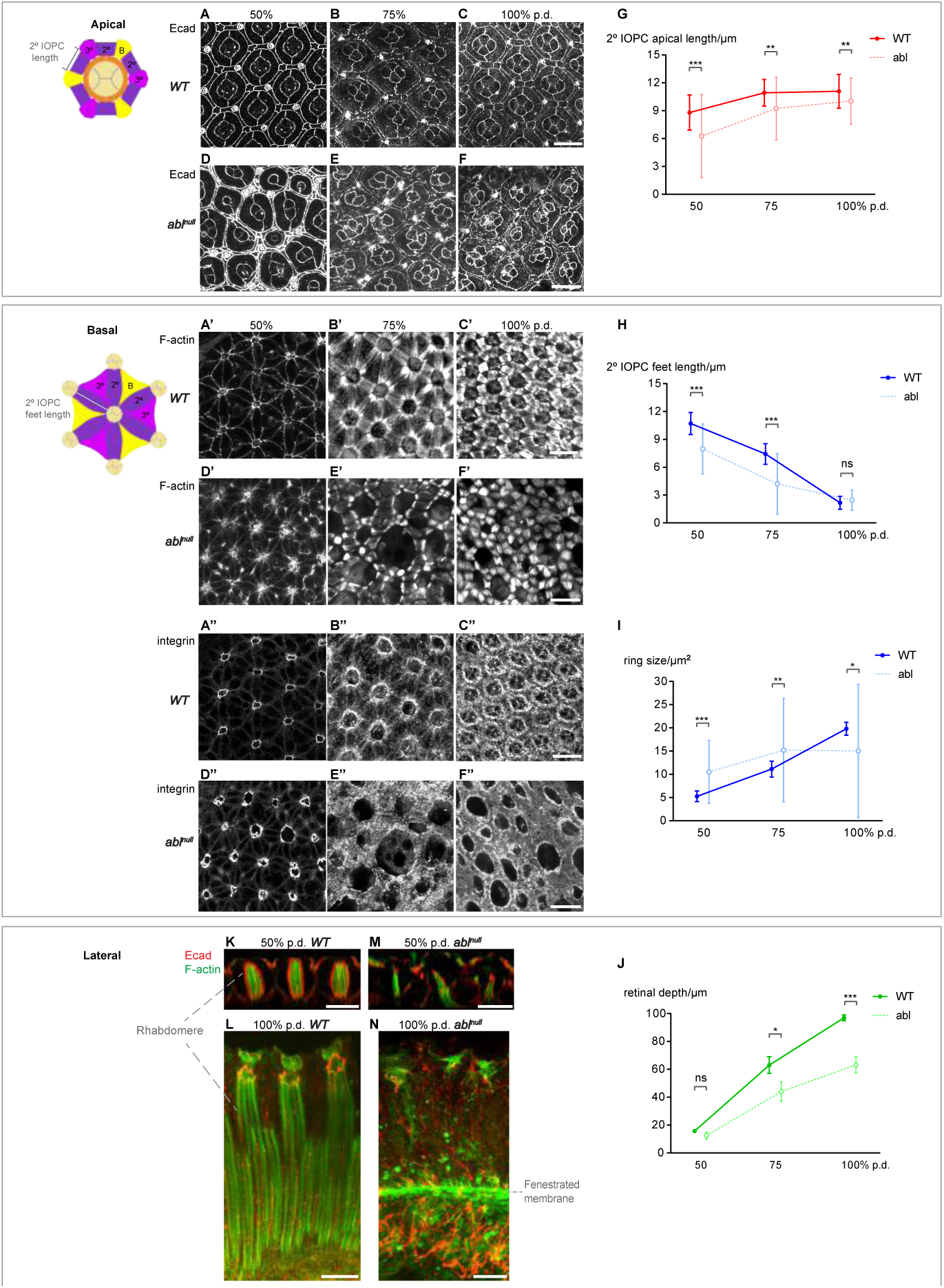
Abl-mediated terminal differentiation specializes the cytoskeletal and junctional structures of the apical, basal and longitudinal networks. **(A-C)** The stereotyped hexagonal pattern of the WT apical network, with single ommatidium schematic color-coded as in Figure 1B. **(D-F)** Abl loss disrupts the apical network, although pattern improves over time. **(A’-C’)** Elaboration and isotropic contraction of the WT basal network, with schematic. **(D’-F’)** Abl loss disrupts the basal network, precluding its isotropic contraction. **(A”-C”)** Attachment points of the basal network to the central rings are reenforced by anchorage to the underlying ECM. **(D”-F”)** Abl loss disrupts basal network-ECM connections, contributing to loss of pattern and integrity. Note that in (**A-F”)**, for each time point and genotype, the same disc, and the same set of ommatidia, were imaged at apical vs. basal planes. Scale bars = 10µm. **(G-I)** Plots of mean ± SEM of secondary IOPC apical domain length, secondary IOPC basal foot length and basal ring area. For each time point and genotype, measurements were made in 30 ommatidia/retina, n= minimum of 4 retinas. **(J)** Plot of mean ± SEM of retinal depth. For each time point and genotype, measurements were made in the central-most 10 ommatidia/retina, n= 3 retinas. **(K,L)** Lateral reconstructions of WT retinas showing the establishment and elaboration/elongation of the longitudinal network of rhabdomere bundles. **(M,N)** Lateral reconstructions of abl^null^ retinas show the collapse of the longitudinal network, and the associated defects in retinal elongation and integrity. Scale bars = 10µm.

In contrast, the shapes, positions and contacts of *abl^null^* retinal cells were aberrant, showed heterogeneity within the same cell type and changed over the 90min course (Figure 1G; Figure S2C and S2D). Most photoreceptor cell bodies had dropped basally and rhabdomeres appeared disorganized. Occasional photoreceptors were detected breaching the fenestrated membrane (Figure 1G, cyan cells) leaving uneven spacing between neighboring ommatidia. The surrounding IOPCs rearranged their position and shape to accommodate the falling photoreceptors, thereby keeping the remaining epithelial sheet intact (Figure 1G and Video S2). The resulting irregular apical and basal contacts impacted pattern along both surfaces (Figure S2C and S2D). Together, these observations suggest that the distinct shapes and spatial arrangement of the photoreceptors and IOPCs is critical to the tissue’s ability to withstand morphogenetic change and maintain integrity.

### Abl is required to elaborate the specialized cytoskeletal domains of both photoreceptors and IOPCs that together organize 3D tissue pattern

To study how morphogenetic remodeling in photoreceptors and IOPCs collectively maintains retinal 3D organization, we examined the consequences of Abl loss to each cell type and to tissue patterning along the different epithelial planes (Figure 2). To bridge cellular and tissue scale analyses, we further conceptualized the specialized cytoskeletal domains that organize the apical, basal and longitudinal planes as a 3D structural scaffold (Figure 1F’). IOPC apical and basal domains provide hexagonally patterned in-plane connections while the photoreceptor rhabdomeres mediate out-of-plane coupling, physically bridging the apical and basal surfaces through anchorage to the cone cell caps and feet embedded in each plane. Scaffold alignment and connections are maintained throughout the elongation phase despite extensive remodeling of the cellular structures. We hypothesized that structural integrity and correct organization of the 3D scaffold is crucial to maintain retinal integrity. Therefore, defects in the terminal differentiation programs that establish the cellular structures and connections that define the scaffold should contribute to the collapse of *abl^null^* photoreceptors. As they fall, the photoreceptors may further perturb scaffold structures and connections.

To identify the cellular defects that disrupt tissue-level pattern, we compared wildtype and *abl^null^* retinas along each plane from 50% p.d. when photoreceptor falling begins to 100% p.d. when the final adult form is achieved. We used F-actin to highlight the specialized cytoskeletal structures that give retinal cells their distinct shapes and Ecadherin (Ecad) to mark the adherens junction connections that organize them into ommatidial units. Focusing first on the apical network, in *abl* mutant retinas, variable apical cell shapes, sizes and cell-cell contacts at 50% p.d. precluded regular hexagonal packing and suggested an anisotropic tension distribution (Figure 2D vs. 2A). For example, multiple rows of IOPCs, rather than a single secondary IOPC, separated adjacent ommatidia while some cone cells dropped sub-apically, disrupting the stereotyped quartet-pattern of apical contacts (Figure 2D). By 100% p.d., apical network pattern was improved, as indicated by the more regular ommatidial shapes and IOPC and cone cell apical profiles (Figure 2E and 2F vs, 2B and 2C). Measurements of secondary IOPC length as a proxy for the regularity of hexagonal pattern confirmed these observations (Figure 2G). The gradual improvement in regularity of ommatidial hexagonal packing suggested that IOPCs continuously optimize their contacts within the apical network, allowing them to recover a more uniform tension distribution after the disturbance of the falling photoreceptors moved away from the apical toward the basal side.

An opposite temporal progression was observed in the basal network (Figure 2A’-2F’’). At 50% p.d., F-actin localization outlined a recognizable radial pattern of IOPC feet in *abl^null^* retinas (Figure 2D’ vs. 2A’) although heterogeneity in the central rings suggested tension was unevenly distributed across the plane (Figure 2D” vs. 2A”). Over time, although the IOPC footprints contracted as in wildtype (Figure 2H), the radial alignment of their F-actin bundles, their connections to the central rings, and the underlying ECM all appeared increasingly disorganized and variable (Figure 2E’-2F” vs. 2B’-2C”; 2I). The increasing severity of basal network disruption over time paralleled the temporal progression of photoreceptor collapse and loss (Figure 1), suggesting that in response to the physical damage caused by the falling photoreceptors, the IOPC feet rearranged their cell-cell and cell-ECM contacts to fill the tissue space and minimize loss of integrity.

The disorganization and heterogeneity within the apical and basal networks predicted the two tissue planes would be out of register. We examined the longitudinal cytoskeletal network that normally provides this alignment. The disruptions of the photoreceptor rhabdomeres caused by *abl* loss were striking (Figure 2H and 2K-2N). At 50% p.d., wildtype rhabdomere clusters align perfectly along the longitudinal axis, span the full tissue depth with anchor points at the apical cone cell caps and basal feet, and their cytoskeletal domains are surrounded by an adherens junction belt that connects neighboring photoreceptors within each ommatidium (Figure 2K). In contrast, in *abl* mutant retinas, cytoskeletal domains and adhesions were fragmented and were no longer oriented orthogonal to the apical and basal surfaces. Junctional connections to the overlying cone cells appeared disrupted and basal collapse was evident (Figure 2M). By 100% p.d., the fragmentation and misalignment of *abl* mutant rhabdomeres (Figure 2N) was in sharp contrast to the intact well-aligned rhabdomere bundles of wildtype (Figure 2L). Below the fenestrated membrane, a tangled mass of Ecad-marked apical membrane reported the collapse of *abl* mutant photoreceptors through the retinal floor (Figure 2N). Retinal depth was significantly reduced relative to wildtype (Figure 2N vs. 2L; Figure 2J). Together these phenotypes suggest Abl function is required at the cellular level to produce the specialized shapes, structures and connections of the photoreceptors and IOPCs. At the tissue level, the structural integrity of the entire 3D scaffold relies on these cellular features; if they are disrupted, the retinal epithelium cannot complete the morphogenetic program.

### Abl is enriched in the cytoskeletal specializations of both photoreceptors and IOPCs

To define the subcellular contexts for Abl function, we examined Abl protein localization using an endogenously GFP-tagged allele (Nagarkar-Jaiswal et al., 2015). Abl expression was detected in all retinal cell types, with enrichment in the IOPC and photoreceptor cytoskeletal and junctional domains that define the 3D structural scaffold (Figure 3 and Figure S3). For example, Abl overlapped with F-actin along the full length of the developing (Figure 3A-3D and Figure S3A and S3B) and mature rhabdomeres (Figure S3D and S3E). Abl was also detected in the IOPCs (Figure 3B,3B’ and 3D,3D’) where the overlap with F-actin was particularly striking in the basal feet (Figure 3D,3D’ and Figure S3). The expression and localization of Abl in both the photoreceptors and IOPCs position it in time and space to regulate the specialization of the F-actin based scaffold.

**Figure 3.**
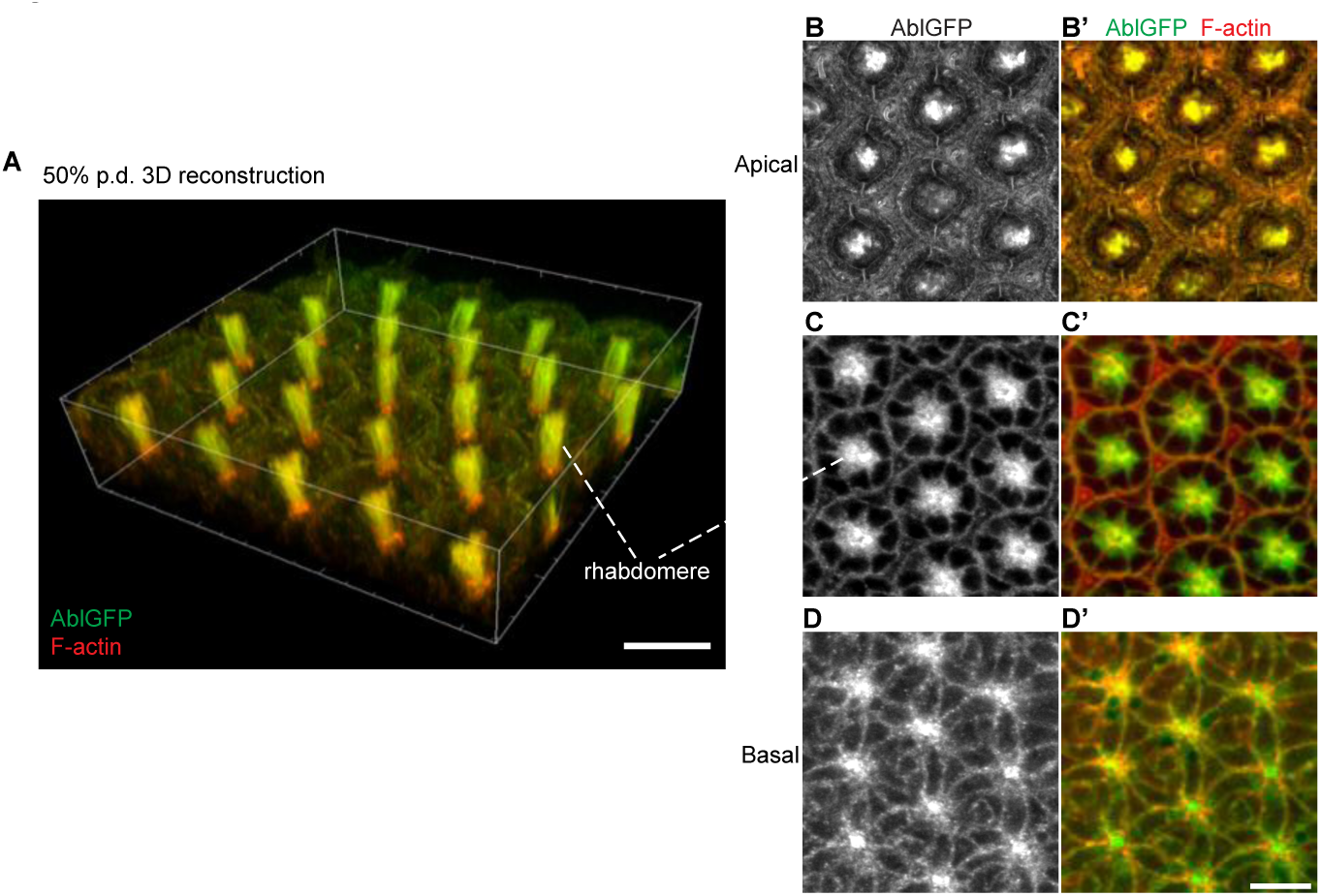
Abl is enriched in the photoreceptor and IOPC F-actin networks along all planes of the 3D scaffold. **(A)** 3D reconstruction showing Abl^mimicGFP^ enrichment in the F-actin-rich longitudinal network of a WT 50% p.d. retina. Sections encompassing the most apical and basal planes shown in **B** and **D** were omitted for clarity. **(B,B’)** Apically, Abl^mimicGFP^ enrichment is strongest at the rhabdomere-cone cell anchor points in the center of each ommatidium, with lower levels detected in cone cells and IOPCs, but not in primary PCs. **(C,C’)** A subapical plane (dashed line) shows Abl^mimicGFP^ enrichment in rhabdomeres. **(D,D’)** Basally, Abl^mimicGFP^ overlaps F-actin at the rhabdomere-cone cell feet anchor points and outlines the basal network of IOPC feet. Scale bars = 10µm.

### Abl has Ena-dependent functions in the photoreceptors and Ena-independent functions in the IOPCs

Assembly of branched F-actin networks supports the specialized structures in both photoreceptors and IOPCs (Galy et al., 2011; Signore et al., 2018). Given that a balance between linear and branched F-actin networks is required for a cell to build specific cytoskeletal structures (Burke et al., 2014; Kannan et al., 2017; Suarez and Kovar, 2016), and that Enabled (Ena) promotes linear F-actin assembly and is inhibited by Abl in a variety of cellular contexts (Comer et al., 1998; Forsthoefel et al., 2005; Fox and Peifer, 2007; Gates et al., 2007; Gertler et al., 1995; Grevengoed et al., 2001; Kannan et al., 2014; Kannan et al., 2017; Lin et al., 2009; Rogers et al., 2021), we asked if the mechanisms underlying Abl function in the photoreceptors and IOPCs involved negative regulation of Ena. Using rhabdomere organization (Figure 4A-4B’) and photoreceptor nuclear position (Figure 4C-4D’) at the subapical plane as readouts, we found that heterozygosity for *ena*, which on its own did not perturb retinal development, suppressed the *abl* mutant phenotype. This result could reflect Abl inhibition of Ena in the photoreceptors, in the IOPCs or in both cell types. We examined Ena expression to distinguish between these possibilities. Overlap with F-actin was detected in wildtype photoreceptor rhabdomeres (Figure 4E and 4E’) whereas in *abl^null^* ommatidia Ena subcellular localization was disrupted and appeared more basally dispersed (Figure 4G and 4G’). Within the IOPCs, low Ena levels were detected in their apical domains (Figure 4E and 4E’) but not in their basal feet (Figure 4F and F’). These expression differences predicted that Abl deploys Ena-dependent regulation in the photoreceptor rhabdomeres but acts via Ena-independent mechanisms in the IOPC contractile feet.

**Figure 4.**
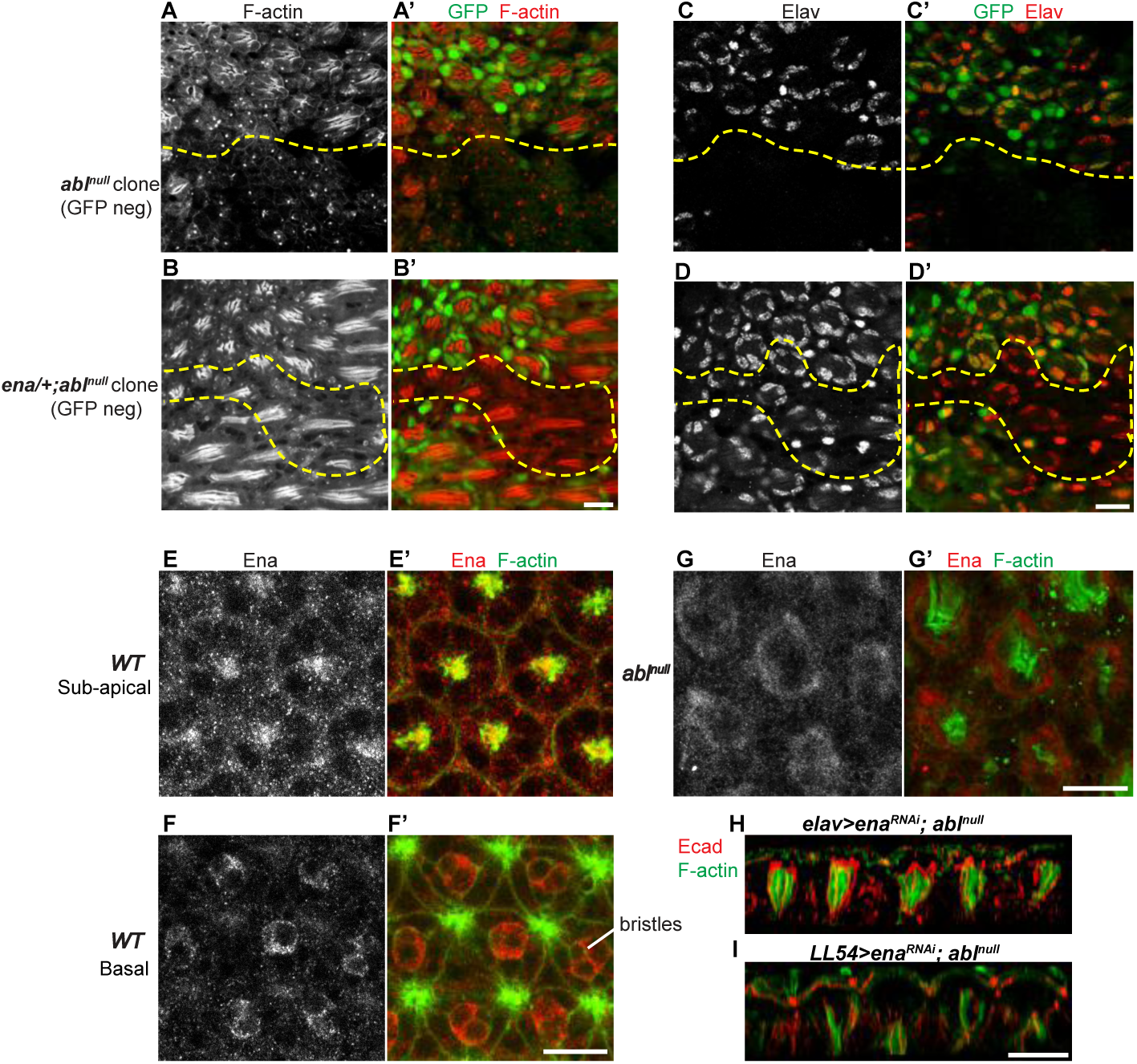
Abl uses Ena-dependent and independent mechanisms to regulate the cytoskeleton in photoreceptors vs. IOPCs. **(A-D’)** Subapical planes of 75% p.d. retinas with GFP-negative *abl^null^* (*abl^2^)* clones, with clonal boundaries marked by yellow dashed lines, show that reducing *ena* dose suppresses the terminal differentiation defects associated with photoreceptor “falling”. **(E,E’)** Subapical plane showing Ena localization to the rhabdomeres and photoreceptor cell bodies. **(F,F’)** Basal plane showing Ena enrichment in bristles, but not in the IOPC feet. **(G,G’)** Subapical plane showing loss of rhabdomeric Ena enrichment in *abl^null^*. **(H,I.)** Lateral plane reconstructions at 50% p.d. showing that photoreceptor-specific Ena knockdown, but not IOPC-specific Ena knockdown, improves rhabdomere structure, organization and spacing in *abl^null^* retinas. Scale bars = 10µm.

To test this, we selectively expressed Ena dsRNA (Ena^RNAi^) in either photoreceptors or IOPCs in an *abl^null^* background. Photoreceptor-specific Ena^RNAi^ improved the orientation, regularity and organization of the rhabdomeres (Figure 4H) whereas IOPC-specific Ena^RNAi^ did not (Figure 4I). Restoring an organized rhabdomere core to each ommatidium (Figure 4H) also significantly improved surrounding IOPC planar pattern in a non-autonomous manner (Figure S4). These results suggest that the terminal differentiation programs of the photoreceptors and IOPCs rely on distinct Abl-mediated regulation to organize the specialized F-actin structures that underlie their unique cellular morphologies.

### Local interactions between photoreceptors and IOPCs organize the 3D scaffold

Our results above suggested that the distinctive terminal differentiation programs of photoreceptors and IOPCs collectively elaborate a 3D scaffold that maintains tissue organization and integrity. The observation that proper rhabdomere organization could non-autonomously correct IOPC patterning defects (Figure 4H) raised the possibility that interactions between these two retinal cell types coordinate this process. We considered two possibilities. First, one cell type might be the primary organizer, with their shape, position and connections imposing dynamic physical constraints that influence how the terminal differentiation program of the other unfolds. Alternatively, there might be redundancy, with local interactions between photoreceptors and IOPCs mutually constraining and influencing the structures and contacts they each contribute to the scaffold.

As we had shown that Abl is required for both photoreceptor and IOPC terminal differentiation, we tested these possibilities by restoring Abl function selectively to one cell type and then assessing the impact on the other *abl* mutant cell type and on overall retinal organization. We used the partial rescue strategy instead of a selective knockdown approach because *abl^RNAi^* does not recapitulate *abl^null^* phenotypes in the retina (Xiong et al., 2009). Control experiments confirmed the specificity of the genetic strategy, with the expected restriction of *elav-Gal4* and *LL54-Gal4* driven expression to photoreceptor and IOPCs, respectively (Figure S5A-S5F), and no leaky expression or rescue with *UAS-Abl^GFP^* alone (Figure S5G-S5L).

Focusing on 50% p.d., the timepoint marking completion of scaffold establishment, we first asked how restoring Abl to the photoreceptors influenced IOPC shapes and apical and basal network patterns in an otherwise *abl^null^* retina (elav>Abl, abl^null^). As expected, expressing Abl specifically in the photoreceptors restored their morphology, with marked improvement in organization and alignment of the rhabdomere bundles that pattern the longitudinal network, and prevented their basal collapse (Figure 5A–5C). When we examined the IOPC response along the apical plane, their apical profiles revealed significant improvement in the regularity of apical network pattern (Figure 5E-5G and 5M). Non-autonomous rescue of basal network pattern was also evident, with reduced variation in ring sizes indicating a more uniform distribution of tension (Figure 5I-5K and 5N). These results suggested that correct photoreceptor cell morphology, and by extension an intact longitudinal network, promotes correct IOPC morphology and pattern within the apical and basal networks.

**Figure 5.**
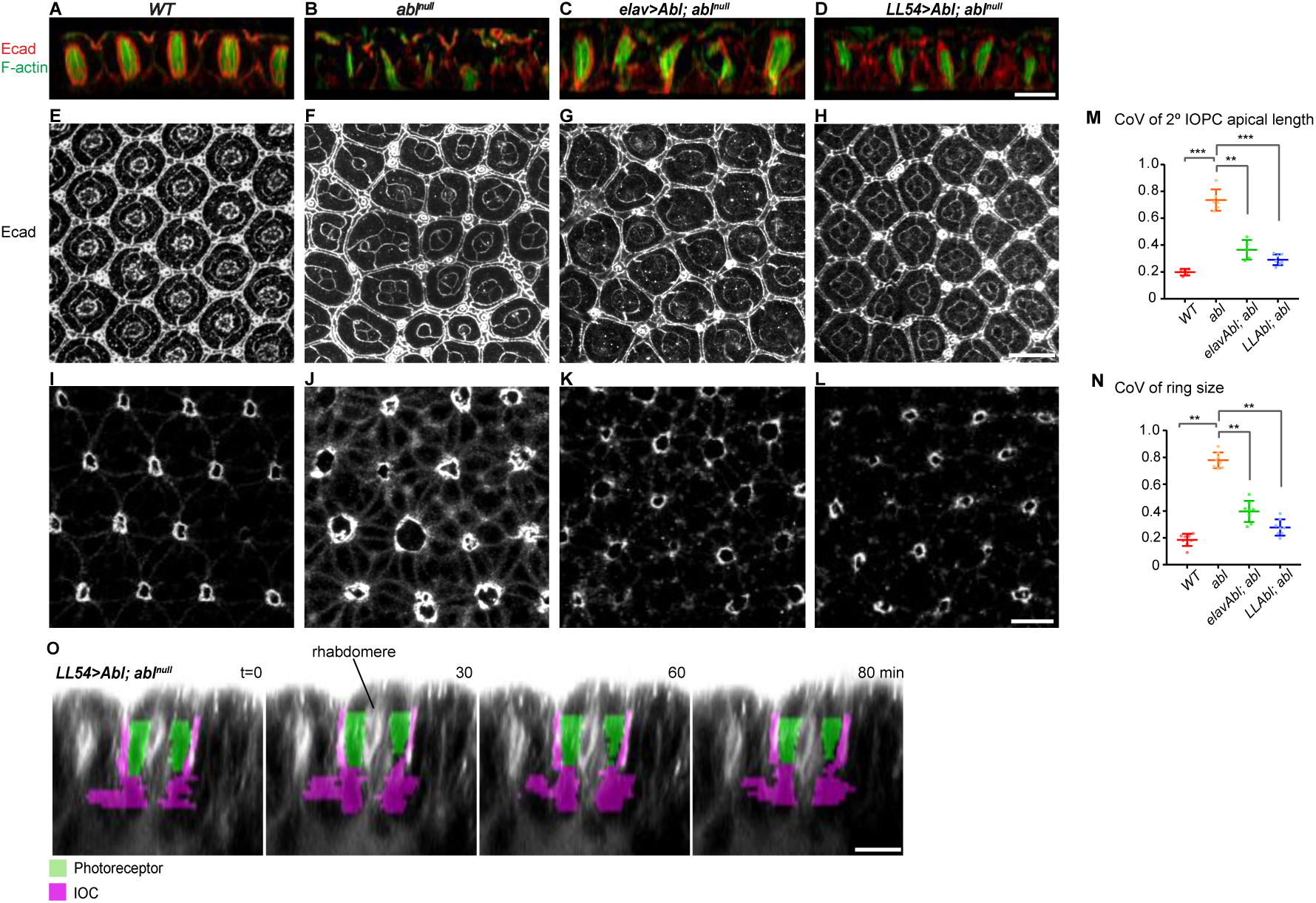
Cell-type specific rescue experiments reveal that local interactions between photoreceptors and IOPCs coordinate 3D scaffold organization. **(A-D)** Lateral, **(E-H)** apical and **(I-L)** basal views of 50% p.d. retinas highlight the sufficiency of restoring Abl function either to the photoreceptors or to the IOPCs in an otherwise *abl^null^* retina to organize the apical, longitudinal and basal network cytoskeletal structures and junctional connections into an intact 3D scaffold. Scale bars = 10µm. **(M,N)** Plots of the CoV of apical secondary IOPC length and of basal ring size. For each genotype, and for each data point, measurements were made in at least 30 ommatidia/retina, n= minimum of 4 retinas. **(O)** False colored stills from a time-lapse movie (see Video S3) of a 50% p.d retina shows that selective restoration of Abl to the IOPCs rescues IOPC and photoreceptor cell shapes, rhabdomere integrity and 3D tissue organization. Scale bar = 10µm.

Second, in the reciprocal experiment we examined the response of *abl^null^* photoreceptors to IOPC-specific restoration of Abl (LL54>Abl, abl^null^). As expected, expressing Abl in the IOPCs rescued their cellular morphology and improved apical (Figure 5H and 5M) and basal (Figure 5L and 5N) network patterns. When we examined the photoreceptor response in the longitudinal plane, the organization, alignment and apical/basal junctional connections of the rhabdomere bundles were all improved, indicating suppression of their basal collapse (Figure 5D, compare to 5B and 5A). Time-lapse imaging further emphasized how the improved overall 3D organization prevented the photoreceptor clusters from collapsing basally; overall, the characteristic shapes and relative positions of both cell types resembled those of a wildtype retina (Figure 5O, Figure S6A and S6B and Video S3). Together, these partial rescue experiments suggested that interactions between these two major retinal cell types, with each non-autonomously influencing the morphology of the other, redundantly coordinate the organization, orientation and integrity of the 3D scaffold.

### Mechanical feedback within the 3D scaffold coordinates terminal differentiation programs in different cells to maintain robust tissue organization

The non-autonomous influence of correct cellular morphology and pattern along an individual plane on cellular morphology and pattern along the orthogonal plane suggested that interactions between the IOPCs and the photoreceptors are an integral organizing feature of the 3D scaffold. These interactions could be primarily passive, with cell shapes adjusting to fill available tissue space, or more active, with morphogenetic change in each cell type promoting the terminal differentiation program of the other. Given the extensive deposition of cellular material and elaboration of structure that occurs within the longitudinal and basal networks during elongation, we predicted that active interactions would be needed to coordinate the completion of these two terminal differentiation programs.

To test this, we used the same cell-type specific partial rescue strategy and assessed rhabdomere expansion, fenestrated membrane contraction, tissue elongation and epithelial integrity in 100% p.d. retinas. We first asked whether selective restoration of Abl function to the photoreceptors (elav>Abl, abl^null^), which supported the elaboration and elongation of a mature longitudinal network of rhabdomere bundles (Figure 6I), was sufficient to rescue the structure and function of the basal network of contractile IOPC feet. In contrast to the disorganized pattern seen in *abl^null^* (Figure 6B), in elav>Abl, abl^null^ partial rescue retinas, the radial alignment of actin bundles in the IOPC feet and their connections to the central rings appeared more uniform (Figure 6C vs. 6A), suggesting successful elaboration of the cytoskeletal structures and connections needed for isotropic contraction of the fenestrated membrane. Similar results were obtained by genetically suppressing the *abl^null^* phenotype with photoreceptor-specific Ena knockdown (elav>Ena^RNAi^, abl^null^) while little or no suppression was obtained with IOPC-specific Ena knockdown (LL54>Ena^RNAi^, abl^null^), consistent with Abl using an Ena-dependent mechanism in the photoreceptors and an Ena-independent mechanism in the IOPC basal stress fibers (Figure 6E and 6F). Measurement of ring size, a metric that summarizes IOPC shape, radial actin alignment and regularity of tissue-level basal network pattern (Figure 6G and 6H) and of retinal depth (Figure 6K) confirmed rescue of the entire tissue-level morphogenetic program and maintenance of retinal integrity. These results suggested that photoreceptor-IOPC interactions actively induce morphogenetic remodeling in the IOPCs, and that at the tissue level, an intact longitudinal network can nonautonomously rescue the basal network, thereby reconstructing an intact scaffold capable of maintaining retinal integrity during elongation.

**Figure 6.**
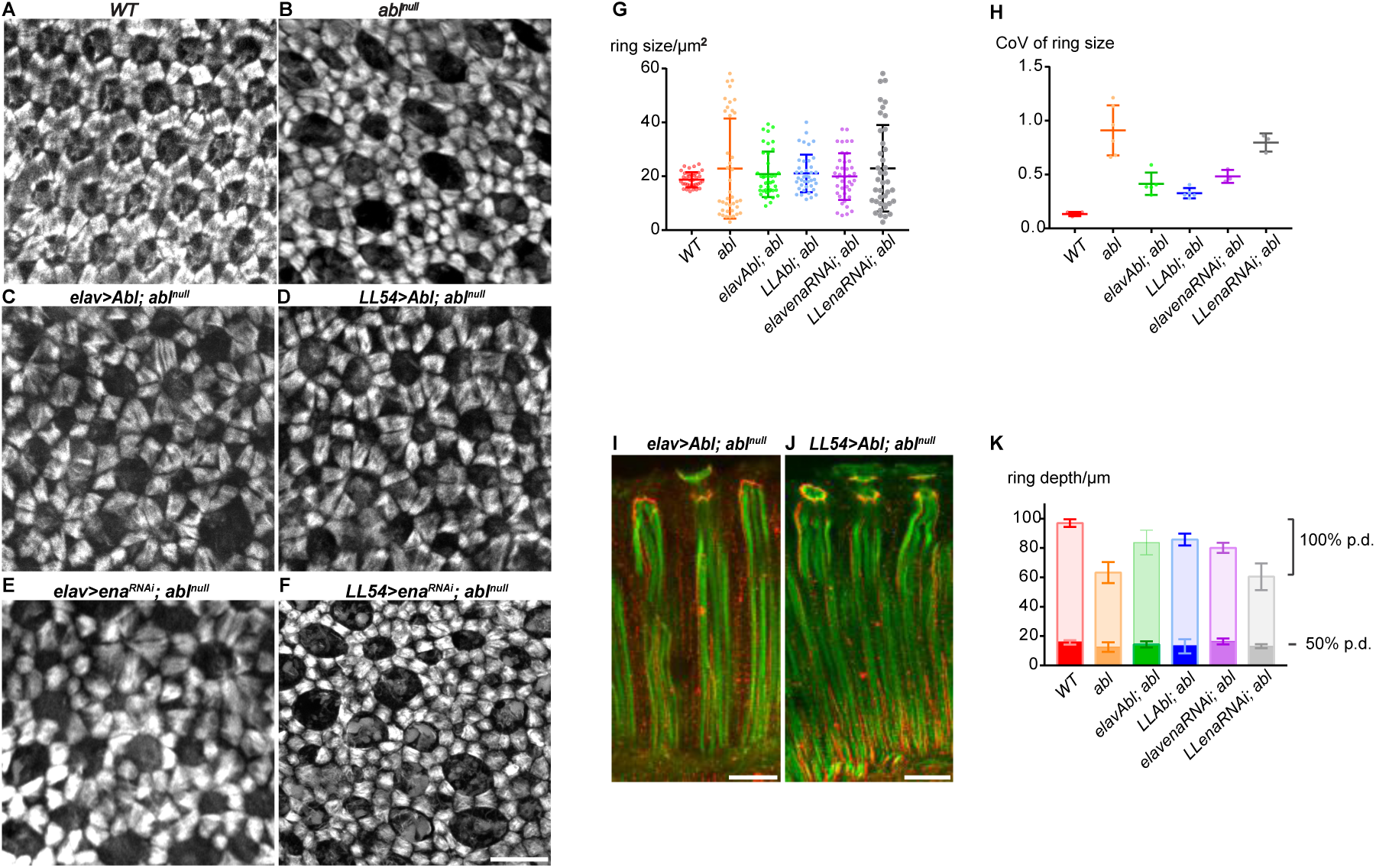
Feedback within the 3D scaffold coordinates photoreceptor and IOPC terminal differentiation programs to maintain tissue organization and integrity during retinal elongation. **(A-F)** Basal F-actin patterns in the IOPC feet of 100% p.d. retinas highlight the sufficiency of a correct photoreceptor terminal differentiation program to non-autonomously induce a basal network with proper radial orientation, symmetry and regularity of IOPC feet in an otherwise *abl^null^* retina. Scale bars = 10 µm. **(G)** Plot showing mean ± SEM of 100% p.d. ring size. For each genotype, data show measurements made in at least 30 ommatidia from a single retina. **(H)** Plot of CoV of ring size. For each genotype, and for each data point, measurements were made in at least 30 ommatidia/retina, n= minimum of 4 retinas. **(I,J)** Lateral views of 100% p.d. rhabdomeres highlight the sufficiency of IOPC-specific expression of Abl to non-autonomously induce active remodeling of photoreceptor rhabdomeres and tissue elongation in an otherwise *abl^null^* retina. Scale bars = 10 µm. **K.** Plot showing mean ± SEM of retinal depth at 50% (dark colored bars) and 100% (light colored bars) p.d.. For each time point and genotype, measurements were made in the central-most 10 ommatidia/retina, n= 3 retinas.

Finally, we examined the reciprocal partial rescue experiment to ask whether elaboration of correct apical/basal network pattern was sufficient to induce remodeling and elongation of the photoreceptor rhabdomeres (LL54>Abl, abl^null^). As expected, restoring Abl function to the IOPCs resulted in correct elaboration of the basal network (Figure 6D, 6G and 6H). Remarkably, this non-autonomously rescued the terminal differentiation program of the *abl^null^* photoreceptors such that they elaborated well-organized and properly oriented rhabdomeres that spanned the full longitudinal axis of the epithelium (Figure 6J) and maintained junctional attachments to the apical cone cell caps (Figure 6J) and basal cone cell feet (Figure S6C-E). Not only were rhabdomere structures, junctional contacts and organization restored, but their elongation was also recovered (Figure 6K), indicating active remodeling of this highly specialized structure. We conclude that interactions between photoreceptors and IOPCs, mediated through the 3D scaffold structures and connections, actively coordinate their terminal differentiation programs, and that together this confers physical robustness to the tissue-level morphogenetic program.

## Discussion

Organ form and function derives from the precise arrangement of different cell types with various sizes, shapes and specialized structures. In complex tissues, coordinating different cellular morphogenetic events in 3D to achieve the correct final form presents a major developmental challenge. Here we showed that the Drosophila pupal retina resolves this challenge by forming a physically coupled 3D cytoskeletal scaffold that integrates two modules of regulation: cell-type specific terminal differentiation programs mediated by Abl specialize the cytoskeletal domains that pattern the individual planes of the scaffold; and tissue-intrinsic feedback relays within the scaffold coordinate morphogenetic change between different retinal cell types. Both modules together ensure the fidelity and integrity of retinal morphogenesis. We propose such 3D scaffolds might be a general feature of tissue morphogenesis, taking on context-specific forms but providing analogous cell and tissue-level coordination to the acquisition of 3D organ form.

### Mechanical feedback matches developmental progress in different cells to coordinate 3D tissue morphogenesis

Late pupal retinal morphogenesis involves drastic shape changes at the cell and tissue level. In contrast to many epithelia where tissue elongation and growth involves changes of junctional contacts through cell intercalation, division and motility along the axes of elongation/growth (Aigouy et al., 2010; Baena-López et al., 2005; Blankenship et al., 2006; Clarke and Martin, 2021; Dye et al., 2017; Dye et al., 2021; Etournay et al., 2015; Glickman et al., 2003; Irvine and Wieschaus, 1994; Mao et al., 2013; Paré and Zallen, 2020; Shindo, 2018; Silva and Vincent, 2007; Warga and Kimmel, 1990; Wilson and Keller, 1991; Zallen and Wieschaus, 2004), the pupal retina relies on a persisting network of cell-cell contacts and cytoskeletal domains to generate and withstand the morphogenetic changes during elongation that produce adult organ form.

One significant finding in this study is that photoreceptors and IOPCs coordinate their morphogenetic programs in order to reenforce tissue organization during morphogenesis (Figure 7A). First, consistent with previous studies (Baumann, 2004; Longley and Ready, 1995; Ready, 2002), our 3D reconstructions highlight the concomitant elaboration of photoreceptor rhabdomeres with the elaboration and contraction of IOPC feet. As both cytoskeletal structures converge from different planes to anchor to the cone cell feet, the cellular coordination between photoreceptors and IOPCs stems from the physical coupling between their specialized cytoskeletal structures along orthogonal planes. Second, restoring Abl function in either the photoreceptors or IOPCs is sufficient to support the terminal differentiation program of the other *abl* mutant cell type. This result shows that the non-autonomous rescue is not merely a passive restoration of cell position and shape, but also induces active remodeling of the cytoskeletal domains and structural specializations through an Abl-independent mechanoresponse. Strikingly, when Abl was specifically restored to the IOPCs, the refined planar pattern not only provided physical support that kept the photoreceptors from falling out of the epithelium, but also induced elaboration and expansion of rhabdomeres in the absence of Abl. On the other hand, rescuing the organization of the central photoreceptor cell clusters by restoring Abl or reducing Ena dosage in photoreceptors was sufficient not only to correct the position and shape of the surrounding IOPCs, but also to direct the radial assembly of contractile IOPC feet. In both scenarios, the reiteration of local interactions between photoreceptors and IOPCs across the ommatidial field restored large-scale tissue order, implying an underlying tissue-level orthogonal coupling that coordinates diverse morphogenetic events in 3D space to achieve robust retinal morphogenesis.

**Figure 7.**
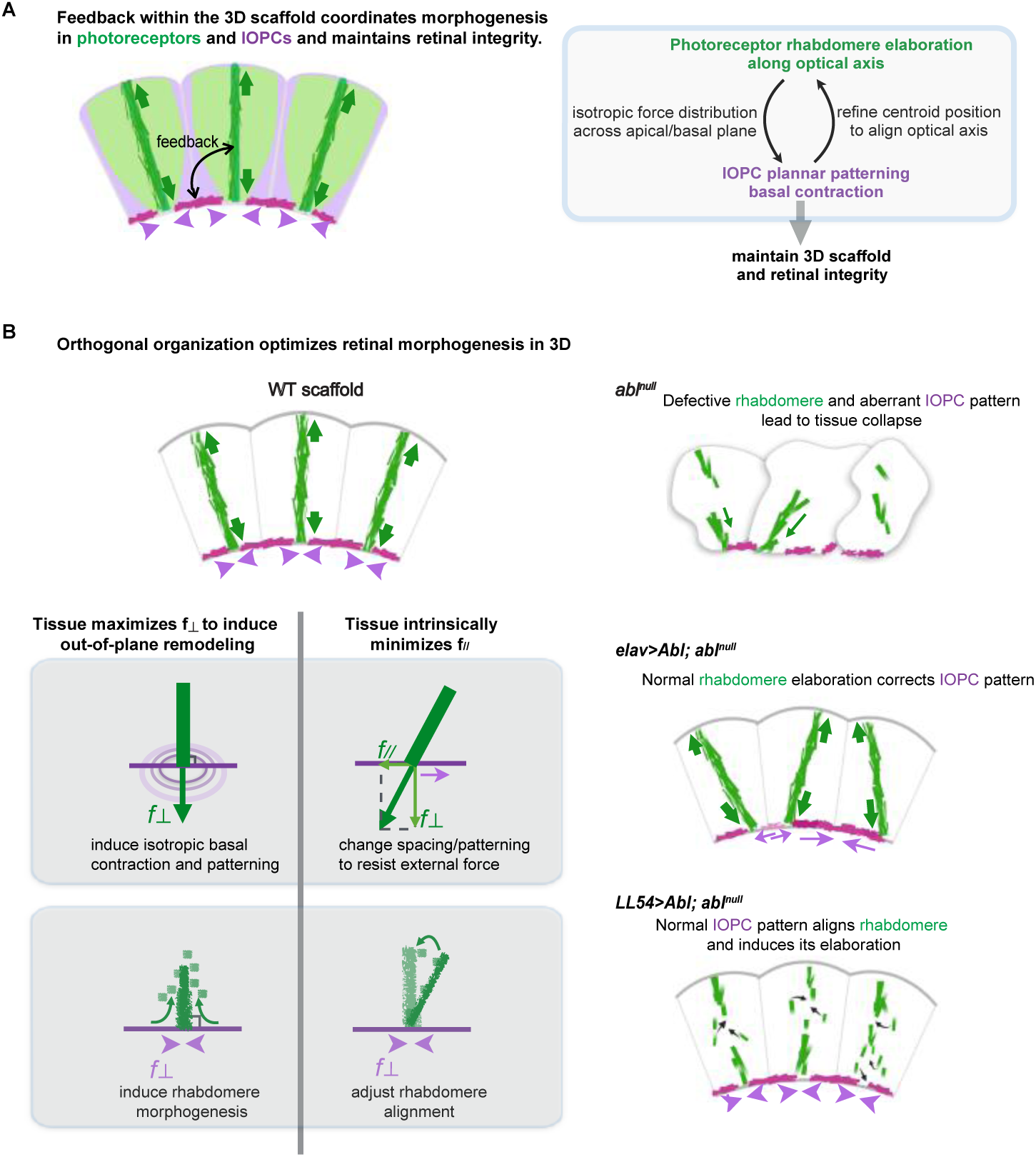
Model of feedback within the 3D scaffold. **(A)** Schematic depicting the spatial organization of photoreceptors (light green) and IOPCs (light pink). Black arrow represents feedback between them as they elaborate their specialized cytoskeletal structures (dark green, elongating rhabdomeres; dark pink, contractile basal feet). Green arrows depict the force exerted by rhabdomere elongation along the optical axis. Pink arrowheads depict the isotropic contraction of the retinal floor in each ommatidium. **(B)** Schematics showing how the orthogonal structure of the 3D scaffold actively communicates and coordinates concurrent morphogenetic changes in different cell types across different tissue axes. Pink arrows depict the response of the retinal floor to external forces exerted by the rhabdomeres.

### Mechanical feedback through an orthogonally coupled 3D scaffold mediates cellular communication

One way to explain this tissue-level coordination is through feedback within the mechanically coupled 3D cytoskeletal adhesion network architecture. Previous studies described planar supracellular networks that span multiple cell apices as central to many morphogenetic processes including epithelial closure, elongation, invagination and folding (Kiehart et al., 2000; Martin et al., 2009; Popkova et al., 2021; Yevick et al., 2019). Our study extends this concept to 3D. We suggest that the specialized cytoskeletal/junctional domains of the photoreceptors and IOPCs provide a mechanically coupled 3D scaffold that channels cell and tissue-scale feedback. Through these interactions, cytoskeletal structure and pattern in one cell type along one plane promotes cytoskeletal organization and pattern in the other cell type along the orthogonal plane. This positive feedback system allows dynamic communication between cells that ensures the concomitancy of their morphogenetic progress. Given the significant morphogenetic changes that accompany the final stages of retinal elongation, incorporating intrinsic feedback between different planes in the 3D scaffold may be critical for maintaining robust tissue organization.

Two general mechanisms could underlie a tissue’s ability to communicate between orthogonal axes. One relies on the incompressibility of the cytoplasm, with competition for cell volume constraining cell shapes and coordinating 3D change (Bagnat et al., 2022; Gelbart et al., 2012; Harmansa et al., 2022; Stooke-Vaughan and Campàs, 2018). Alternatively, forces could be transmitted directly through the dedicated cytoskeletal scaffold, with the geometrical arrangement of the scaffold affecting inter-plane communication (Figure 7B). Our finding that restoring one plane of the 3D scaffold could induce remodeling and reorientation of the orthogonal plane suggests that the tissue simultaneously favors orthogonal organization for efficient morphogenesis and intrinsically corrects and minimizes off-axis pattern. If one axis is tilted, the component forces will be exchanged with the opposing plane, which triggers cytoskeletal remodeling or cellular level rearrangement in the opposing plane as in an adaptive planar supracellular network (Aigouy et al., 2010; Chanet et al., 2017; Coen and Rebocho, 2016; Huang et al., 2018; Khalilgharibi et al., 2019; Mao and Baum, 2015; Rebocho et al., 2017; Shyer et al., 2013; Wyatt et al., 2020; Yevick et al., 2019). The component forces strictly coming from the orthogonal plane induce the constructive remodeling along the opposing plane, thereby mechanically coupling rhabdomere expansion and isotropic contraction of IOC feet.

In the wildtype retinal 3D scaffold, rhabdomeres maintain an almost orthogonal arrangement to the curved surfaces of the IOPC apical and basal domains throughout morphogenesis (Figure 7A and 7B). Given that elaboration of rhabdomere structure requires extensive membrane/cytoskeletal biogenesis and deposition (Raghu et al., 2009), in addition to being important for vision, maintaining the optimal orthogonal orientation has the advantage of minimizing biogenesis during elongation. In an *abl* mutant retina where coupling through the dedicated 3D scaffold collapses, the first mechanism (competition for volume) likely dominates but is insufficient to maintain the orthogonal organization that supports full tissue elongation. In the cell-type specific rescues, restoration of one plane exerts forces to the anchor point, triggering the opposing plane to minimize the component forces along the same orientation. Adjusting both axes relative to each other eventually restores the orthogonal organization of the scaffold, such that all in-plane stresses can be efficiently used for remodeling along the orthogonal axis and can support close to wildtype tissue elongation. We speculate that orthogonally coupled cytoskeletal configurations may represent a general strategy not only for communicating and resolving out of register geometries, but also for optimizing the use of in-plane stresses to direct out-of-plane remodeling and thus coordinate robust 3D morphogenetic change.

### Remodeling of cytoskeleton, junctions and ECM all contribute to 3D scaffold structure and integrity

Our study highlights how the 3D scaffold couples orthogonally oriented morphogenetic events to maintain retinal organization. Although our characterization focused on interactions between photoreceptors and IOPCs, additional interactions, such as anchorage to the cone cells and to the basal ECM, contribute to the structural specialization of scaffold components and maintenance of scaffold integrity. The cone cell quartet provides a hub that physically connects both apical and basal planar hexagonal patterns to the orthogonally oriented rhabdomeres. The cone cell feet also plug the central rings and complete the in-plane tiling of the basal floor by making septate junction connections with the IOPC feet (Banerjee et al., 2008). Thus, as the photoreceptor rhabdomeres mature and the IOPC feet elaborate contractile stress fibers, concomitant remodeling of cone cell shape and junctional connections with both cell types will further strengthen the scaffold. Consistent with this, in *abl* mutants, our observation of increased and irregular ring size implicates elaboration of correct cone cell feet shape and attachments as an integral component of scaffold elaboration. Identification of Neurexin IV, a septate junction component, and *DPax2,* a transcription factor that presumably regulates many aspects of cone cell terminal differentiation, as “photoreceptor falling” mutants (Banerjee et al., 2008; Charlton-Perkins et al., 2017), further supports our model.

The retinal floor is comprised not only of the IOPC basal contractile network but is also constrained by the underlying ECM; close coupling between these two components enables coordinated force transmission across the entire plane. IOPCs form integrin-mediated focal adhesions that anchor the contractile fibers to the underlying ECM. Concomitant with contraction of the IOPC basal network, the ECM shows signs of increasing collegen IV (Vkg-GFP) deposition and increased integrin intensity around the central rings (Figure S2E). This suggests that mechanical coupling between the elaborating ECM layer and the IOPC basal network matches their developmental progress during elaboration of the specialized retinal floor, thereby contributing to its integrity and coordination of cellular and tissue-scale events during elongation. Consistent with this, integrins have long been known to be essential to retinal integrity (Longley and Ready, 1995). The specific contributions of different ECM components remain to be explored.

In addition to the cone cell and ECM components, previous studies have identified several other cytoskeletal/junctional components or regulators whose loss compromises retinal integrity such that photoreceptors fall below the retinal floor. The list includes: Arp2/3 and WAVE/SCAR components (Galy et al., 2011); Cofilin (Pham et al., 2008); DRac1 (Chang and Ready, 2000); Pebbled/Hindsight (Pickup et al., 2002); and Afadin/Canoe (Matsuo et al., 1999). A few on the list have been implicated in Abl function in other developmental contexts (Grevengoed et al., 2003; Kannan and Giniger, 2017; Kannan et al., 2017; Yu and Zallen, 2020), and so could provide insight into mechanisms of Abl function in the retina. More broadly, based on our analysis and interpretation of the *abl* loss of phenotype together with the known function of these gene products in regulating actin cytoskeleton and cell-cell junctions, we suggest that this set of genes likely contributes via a variety of mechanisms to the connectivity and mechanosensitive remodeling of the scaffold structure.

In conclusion, we propose that the mechanical connectedness of the 3D scaffold underlies a feedback mechanism that ensures the concurrent remodeling of the different components and coordinates developmental progress across the tissue. Together these interactions maintain robust tissue organization. Future study of the cellular machineries and mechanics of the 3D scaffold coupled with characterization of tissue-level growth, geometry and developmental trajectories will provide further insight into how cell and tissue-scale processes collectively produce complex 3D organ form.

## Acknowledgements

We thank Ed Munro, Ellie Heckscher and Rick Fehon for experimental advice and comments on the manuscript, Saman Tabatabaee for assistance with graphics, Christine Labno and Audrey Williams for live image processing advice, past and present Rebay lab members for helpful discussions and G. Bashaw, S.L. Zipursky, R.W. Carthew and S. Horne-Badovinac for fly stocks. X.S. was supported by NIH R01 EY021459 to I.R., J.D. was supported by NIH T32 GM007183, and N.S.L. was supported by NIH T32 GM007281 and by NIH R01 EY021459 to I.R..

## Methods

### Drosophila genetics

All crosses were carried out at 25°C in standard laboratory conditions. *w^1118^* (3605), *abl^1^* (3554), *abl^2^* (8565), *Abl^mimicGFP^* (59761), *LL54-Gal4* (5129), *Elav-Gal4* (8765), *ena^23^* (8571), *G-TRACE* (2820) were from the Bloomington Stock Center. UAS-Abl-GFP (O’Donnell and Bashaw, 2013) was a gift from G. Bashaw. *UAS-ena^RNAi^* (106484) was from the Vienna Drosophila Resource Center. *vkg-GFP* (CC00791) was from Flytrap (Buszczak et al., 2007). *Chp^4.5^-Gal4* (Mishra et al., 2013) was a gift from S.L. Zipursky. GMR-wIR^13D^ (Lee and Carthew, 2003) was a gift from R. Carthew.

*w^1118^* was used as wildtype in all experiments. To generate *abl^null^* animals, *abl^2^/TM6B* males were crossed to *abl^1^/TM6B* virgins and *abl^1^/abl^2^* (non-Tubby) were selected. To generate *abl^2^* clones, *abl^2^,FRT80B/TM6B* males were crossed to *ey-FLP; GFP^nls^,FRT80B* virgins. To reduce *ena* dose in *abl^2^* clones, *ena^23^/CyO, abl^2^ FRT80B/TM6B* males were crossed to *ey-FLP; GFP^nls^,FRT80B* virgins. For cell type specific rescue experiments, *abl^2^* was recombined with the *UAS-Abl-GFP* transgenes inserted in 86Fb docking site (O’Donnell and Bashaw, 2013). *abl^2^,UAS-Abl-GFP/TM6B* males were crossed to *LL54 (or Elav)-Gal4; abl^1^/TM6B* virgins. To test Abl-Ena interactions in specific cell types, *UAS-ena^RNAi^; abl^2^/TM6B* males were crossed to *LL54 (or Elav)-Gal4; abl^1^/TM6B* virgins. For lineage tracing, *w,GMR-w.IR*;*+/+;chaoptin-Gal4, abl^2^/TM6B* females were crossed to *w; G-TRACE; abl^1^/TM6B* males. GMR-w.IR reduced pigment deposition, improving imaging. *G-TRACE* = *P{w[+mC]=UAS-RedStinger}4, P{w[+mC]=UAS-FLP.D}JD1, P{w[+mC]=Ubi-p63E(FRT.STOP)Stinger}9F6/CyO*.

For developmental staging, white pre-pupae (0hr APF) were selected and aged at 25°C. 50%, 75% and 100% p.d. animals for dissection were selected by time (48, 72 and 96hrs APF, respectively), confirmed by morphological landmarks prior to dissection (Bainbridge 1981) and by cellular features after dissection.

### Immunostaining

50% p.d. pupal eye discs were dissected in PBS and fixed for 10 min in 4% paraformaldehyde in PBT (PBS with 0.1% Triton X-100). For 75% and 100% p.d. dissections, pupal/adult heads were prefixed for 20 min, retina were dissected and fixed for 10 min, washed three times in PBT, blocked in PNT (PBT+3% normal goat serum) for 1 hr, incubated overnight at 4°C in primary antibodies diluted in PNT, washed three times in PBT, incubated in secondary antibodies diluted in PNT for 6 hours, washed three times in PBT and mounted in 90% glycerol in 0.1M Tris pH 8.0 with 0.5% n-propyl gallate.

In cell type specific rescue experiments in which *UAS-Abl-GFP* was ectopically expressed under the control of *LL54-Gal4* or *Elav-Gal4*, F-actin was stained with AlexaFluor-488 Phalloidin. Although GFP and AlexaFluor-488 are both excited by the same wavelength, the *UAS-Abl-GFP* signal faded during our standard staining protocol and was no longer detected with the confocal settings used to image F-actin. For Supplementary Figure 3, to preserve the UAS-Abl-GFP signal, discs were imaged after only a 2hr incubation with AlexaFluor-568 Phalloidin.

Antibodies were from the Developmental Studies Hybridoma Bank (DSHB): mouse anti-Elav (1:50); mouse anti-Ecad (1:500), mouse anti-βPS integrin (1:100), mouse anti-Ena (1:50). Cy3/Cy5/FITC-conjugated secondaries,1:2000 (Jackson ImmunoResearch). AlexaFluor-488 and AlexaFluor-568 Phalloidin,1:1000 (Fisher).

### Fixed Microscopy and image analysis

Fluorescent images were obtained using a Zeiss LSM880 confocal microscope with Airyscan. Image processing and depth and ring size measurements were performed using Fiji. To obtain the image of different planes, multiple z-stack slices (0.4µm interval) that encompass the region of interest were averaged.

3D reconstructions and lateal views were made in Imaris (Bitplane). For views that include intact rhabdomeres, the 3D reconstructions were rotated and cropped to expose the lateral view of one line of aligned ommatidia. Graphing and statistical analyses were performed in Prism7 (GraphPad). For all graphs, data were plotted as mean ± SEM and statistical differences between conditions were determined with two-tailed unpaired t tests.

To measure retinal depth, we chose the central region of the retina for all genotypes to avoid regional variation. We generated multiple lateral views in Imaris and used the average length of 10 ommatidia in the central region as the retinal depth.

### Live imaging

Staged 50% p.d. pupae were selected and the puparium was dissected away to expose the distal (apical) surface of the retina. After injecting ∼0.5uL of CellMask Deep Red Plasma Membrane Stain (Thermo-Fisher) diluted 1:25 in Schneider’s *Drosophila* medium into the head, the pupae were incubated in a humid chamber at 25°C for 4 hours. Injected pupae were mounted in a cover-glass bottom dish such that the region of the head containing the developing retina was directly in contact with the cover-glass in a thin layer of Halocarbon 700 oil (Halocarbon Products). Pupae were oriented and immobilized using a log of petroleum jelly and the imaging vessel was kept humid with blotting paper soaked in Schneider’s *Drosophila* medium (method adapted from (Hellerman et al., 2015)). Mounted retinas were immediately imaged on a Zeiss LSM 880 laser scanning confocal microscope using the 40x oil-immersion objective on AiryScan mode. Optical slices of 1um were acquired through the entire apical-basal depth of the tissue every 5 minutes. Although we cannot rule out the possibility that our protocol stalls development or negatively impacts the tissue, in a control experiment, injected animals continued to develop and 85% (n=20) eclosed with no obvious morphological defects. Because the dye injection resulted in a huge intensity difference between apical and basal planes, Fiji Top-Hat (Spot Radius = 0.75) was run on the entire 4D image stack to equalize signal and enhance cell outlines (Script by G. Landini, adapted for use on hyperstacks by C. Labno: https://blog.bham.ac.uk/intellimic/g-landini-software/). The modified macro can be accessed on the University of Chicago Microscopy Core’s GitHub: Core’s GitHub (https://github.com/UChicago-Integrated-Light-Microscopy/ImageJ_macros).

Individual cells were annotated by hand in ImageJ using the xy-planes because of the ease in distinguishing cell outlines. After manually correcting drift, lateral projections of the resulting time-lapse images were made using ImageJ software.

## Supplemental Figure Legends

**Figure S1. Lineage tracing to examine maintenance of photoreceptor cell fate and position in the retinal epithelium.**

**(A,B)** Genetic elements of the G-TRACE lineage trace system result in coexpression of GFP and RFP in photoreceptor nuclei: *chp-Gal4* drives photoreceptor-specific expression of RFP and FLP. FLP removes the stop in the actin-stop-GFP cassette, establishing a permanent lineage mark (GFP).

**(C,D)** Lineage-traced photoreceptor nuclei in a WT 100% p.d. retina reside apically and are never detected beneath the fenestrated membrane.

**(E,F)** Lineage-traced photoreceptor nuclei in an *abl^null^* 100% p.d. retina suggest that even though cell position changes, with some nuclei found beneath the fenestrated membrane, cell fate is not changed.

Schematic indicating apical (red) and basal (blue) planes imaged. Scale bars = 10µm.

**Figure S2. Time-lapse images and subsequent basal ECM elaboration to examine apical and basal network pattern.**

**(A-D).** Schematics and stills from time-lapse movies (Videos S1 and S2) of 50% p.d retinas injected with CellMask (white). False color shows photoreceptors (green), fallen photoreceptors (cyan) and secondary IOPCs (magenta) in a representative ommatidium. Abl loss results in irregular photoreceptor and secondary IOPC cell shapes and contacts that change over the time-course. Basally, photoreceptor “falling” is marked by the appearance of the cyan-marked cell between IOPC feet.

Scale bars = 10µm

**(E).** Progression of basal ECM deposition at 50, 75 and 100% p.d., visualized with Viking-GFP (collagen IV), occurs in sync with the contraction of the IOPC feet and expansion of the central rings in a WT retina. Scale bars = 10 um.

**Figure S3. Abl is enriched in photoreceptor and IOPC F-actin structures at 25% and 100% p.d..**

**(A-F’)** Apical, subapical and basal planes showing the enrichment of Abl^mimicGFP^ and F-actin in photoreceptors and IOPCs throughout retinal development, beginning prior to scaffold establishment and continuing in the adult scaffold. Scale bar = 10µm.

**Figure S4. Abl-Ena genetic interactions show that photoreceptors non-autonomously affect IOPC apical network pattern. (A-C)** Apical network pattern of 50% p.d. retinas. Scale bars = 10µm.

**(D)** Plots of the CoV of apical secondary IOPC length. For each genotype, and for each data point, measurements were made in at least 30 ommatidia/retina, n= minimum of 4 retinas.

**Figure S5. Specificity of Elav-and LL54-Gal4 expression and lack of rescue by UAS-Abl^GFP^ in the absence of a Gal4 driver.**

**(A-C’)** Elav-Gal4 drives expression specifically in photoreceptors, with no leaky expression detected in IOPCs or cone cells.

**(D-F’)** LL54-Gal4 drives expression specifically in IOPCs, with no leaky expression detected in photoreceptors or cone cells.

**(G-J)** Without a Gal4 driver, the UAS-Abl^GFP^ transgene does not rescue apical or basal network pattern.

Scale bars = 10µm.

**(K,L)** Measurement and CoV of secondary IOPC apical length and basal ring size. Plots of secondary IOPC apical length show measurements made in at least 30 ommatidia in a single disc for each genotype. In the CoV plots, for each genotype, and for each data point, measurements were made in at least 30 ommatidia/retina, n= minimum of 4 retinas.

**Figure S6. IOPC-specific expression of Abl in an otherwise *abl^null^* retina restores correct cell-cell contacts within the scaffold.**

**(A,B)** Schematics and stills from a time-lapse movie (Videos S3) of a 50% p.d *LL54>Abl;abl^null^* retina injected with CellMask (white) showing restoration of apical and basal pattern. False color shows photoreceptors (green) and secondary IOPCs (magenta) in a representative ommatidium. Compare to Figure S2A-D.

**(C-E).** Basal plane of 100% p.d. retinas, with schematic, shows how restoration of Abl to the IOPCs rescues the anchorage of the rhabdomeres to the cone cell feet, suggesting that non-autonomous induction of rhabdomere remodeling reconstructs the structural integrity of the 3D scaffold. Scale bars = 10 µm.

**Video S1. Time lapse movie of a 50% p.d. WT retina injected with CellMask to show cell outlines and rhabdomeres and with a representative tracked ommatidium false-colored to identify photoreceptors and IOPCs.**

Frames are re-aligned to keep the center of the tracked ommatidium constant. Framer rate: 5 min/frame; Duration: 90 min. Scale bar: 10µm. Related to Figure 1 and S2.

**Video S2. Time lapse movie of a 50% p.d. *Abl^null^* retina injected with CellMask to show cell outlines and rhabdomeres and with a representative tracked ommatidium false-colored to identify photoreceptors and IOPCs.**

Frames are re-aligned to keep the center of the tracked ommatidium constant. Framer rate: 5 min/frame; Duration: 90 min. Scale bar: 10µm. Related to Figure 1 and S2.

**Video S3. Time lapse movie of a 50% p.d. *LL54>Abl^GFP^*;*abl^null^* retina injected with CellMask to show cell outlines and rhabdomeres and with a representative tracked ommatidium false-colored to identify photoreceptors and IOPCs.**

Frames are re-aligned to keep the center of the tracked ommatidium constant. Framer rate: 5 min/frame; Duration: 90 min. Scale bar: 10µm. Related to Figure 5 and S6.

## Notes

### Competing Interest Statement

The authors have declared no competing interest.

## References

1. Aigouy, B., Farhadifar, R., Staple, D. B., Sagner, A., Röper, J.-C., Jülicher, F. and Eaton, S. (2010). Cell Flow Reorients the Axis of Planar Polarity in the Wing Epithelium of Drosophila. Cell 142, 773–786.

2. Baena-López, L. A., Baonza, A. and García-Bellido, A. (2005). The Orientation of Cell Divisions Determines the Shape of Drosophila Organs. Curr Biol 15, 1640–1644.

3. Bagnat, M., Daga, B. and Talia, S. D. (2022). Morphogenetic Roles of Hydrostatic Pressure in Animal Development. Annu Rev Cell Dev Bi 38, 375–394.

4. Banerjee, S., Bainton, R. J., Mayer, N., Beckstead, R. and Bhat, M. A. (2008). Septate junctions are required for ommatidial integrity and blood–eye barrier function in Drosophila. Dev Biol 317, 585–599.

5. Bao, S. and Cagan, R. (2005). Preferential Adhesion Mediated by Hibris and Roughest Regulates Morphogenesis and Patterning in the Drosophila Eye. Dev Cell 8, 925–935.

6. Baumann, O. (2004). Spatial pattern of nonmuscle myosin-II distribution during the development of the Drosophila compound eye and implications for retinal morphogenesis. Dev Biol 269, 519–533.

7. Bennett, R. L. and Hoffmann, F. M. (1992). Increased levels of the Drosophila Abelson tyrosine kinase in nerves and muscles: subcellular localization and mutant phenotypes imply a role in cell-cell interactions. Development 116, 953–966.

8. Blankenship, J. T., Backovic, S. T., Sanny, J. S. P., Weitz, O. and Zallen, J. A. (2006). Multicellular Rosette Formation Links Planar Cell Polarity to Tissue Morphogenesis. Dev Cell 11, 459–470.

9. Burke, T. A., Christensen, J. R., Barone, E., Suarez, C., Sirotkin, V. and Kovar, D. R. (2014). Homeostatic Actin Cytoskeleton Networks Are Regulated by Assembly Factor Competition for Monomers. Curr Biol 24, 579–585.

10. Buszczak, M., Paterno, S., Lighthouse, D., Bachman, J., Planck, J., Owen, S., Skora, A. D., Nystul, T. G., Ohlstein, B., Allen, A., et al. (2007). The Carnegie Protein Trap Library: A Versatile Tool for Drosophila Developmental Studies. Genetics 175, 1505–1531.

11. Cagan, R. L. and Ready, D. F. (1989). The emergence of order in the Drosophila pupal retina. Dev Biol 136, 346–362.

12. Chanet, S., Miller, C. J., Vaishnav, E. D., Ermentrout, B., Davidson, L. A. and Martin, A.C. (2017). Actomyosin meshwork mechanosensing enables tissue shape to orient cell force. Nat Commun 8, 15014.

13. Chang, H.-Y. and Ready, D. F. (2000). Rescue of Photoreceptor Degeneration in Rhodopsin-Null Drosophila Mutants by Activated Rac1. Science 290, 1978–1980.

14. Charlton-Perkins, M. and Cook, T. A. (2010). Chapter Five Building a Fly Eye Terminal Differentiation Events of the Retina, Corneal Lens, and Pigmented Epithelia. Curr Top Dev Biol 93, 129–173.

15. Charlton-Perkins, M. A., Sendler, E. D., Buschbeck, E. K. and Cook, T. A. (2017). Multifunctional glial support by Semper cells in the Drosophila retina. Plos Genet 13, e1006782.

16. Clarke, D. N. and Martin, A. C. (2021). Actin-based force generation and cell adhesion in tissue morphogenesis. Curr Biol 31, R667–R680.

17. Coen, E. and Rebocho, A. B. (2016). Resolving Conflicts: Modeling Genetic Control of Plant Morphogenesis. Dev Cell 38, 579–583.

18. Collinet, C. and Lecuit, T. (2021). Programmed and self-organized flow of information during morphogenesis. Nat Rev Mol Cell Bio 22, 245–265.

19. Comer, A. R., Ahern-Djamali, S. M., Juang, J.-L., Jackson, P. D. and Hoffmann, F. M. (1998). Phosphorylation of Enabled by the Drosophila Abelson Tyrosine Kinase Regulates the In Vivo Function and Protein-Protein Interactions of Enabled. Mol Cell Biol 18, 152–160.

20. Daley, W. P. and Yamada, K. M. (2013). ECM-modulated cellular dynamics as a driving force for tissue morphogenesis. Curr Opin Genet Dev 23, 408–414.

21. Dye, N. A., Popović, M., Spannl, S., Etournay, R., Kainmüller, D., Ghosh, S., Myers, E. W., Jülicher, F. and Eaton, S. (2017). Cell dynamics underlying oriented growth of the Drosophila wing imaginal disc. Development 144, 4406–4421.

22. Dye, N. A., Popović, M., Iyer, K. V., Fuhrmann, J. F., Piscitello-Gómez, R., Eaton, S. and Jülicher, F. (2021). Self-organized patterning of cell morphology via mechanosensitive feedback. Elife 10, e57964.

23. Etournay, R., Popović, M., Merkel, M., Nandi, A., Blasse, C., Aigouy, B., Brandl, H., Myers, G., Salbreux, G., Jülicher, F., et al. (2015). Interplay of cell dynamics and epithelial tension during morphogenesis of the Drosophila pupal wing. Elife 4, e07090.

24. Forsthoefel, D. J., Liebl, E. C., Kolodziej, P. A. and Seeger, M. A. (2005). The Abelson tyrosine kinase, the Trio GEF and Enabled interact with the Netrin receptor Frazzled in Drosophila. Development 132, 1983–1994.

25. Fox, D. T. and Peifer, M. (2007). Abelson kinase (Abl) and RhoGEF2 regulate actin organization during cell constriction in Drosophila. Development 134, 567–578.

26. Galy, A., Schenck, A., Sahin, H. B., Qurashi, A., Sahel, J.-A., Diebold, C. and Giangrande, A. (2011). CYFIP dependent Actin Remodeling controls specific aspects of Drosophila eye morphogenesis. Dev Biol 359, 37–46.

27. Gates, J., Mahaffey, J. P., Rogers, S. L., Emerson, M., Rogers, E. M., Sottile, S. L., Vactor, D. V., Gertler, F. B. and Peifer, M. (2007). Enabled plays key roles in embryonic epithelial morphogenesis in Drosophila. Development 134, 2027–2039.

28. Gelbart, M. A., He, B., Martin, A. C., Thiberge, S. Y., Wieschaus, E. F. and Kaschube,M. (2012). Volume conservation principle involved in cell lengthening and nucleus movement during tissue morphogenesis. Proc National Acad Sci 109, 19298–19303.

29. Gertler, F. B., Comer, A. R., Juang, J. L., Ahern, S. M., Clark, M. J., Liebl, E. C. and Hoffmann, F. M. (1995). enabled, a dosage-sensitive suppressor of mutations in the Drosophila Abl tyrosine kinase, encodes an Abl substrate with SH3 domain-binding properties. Genes Dev 9, 521–533.

30. Glickman, N. S., Kimmel, C. B., Jones, M. A. and Adams, R. J. (2003). Shaping the zebrafish notochord. Development 130, 873–887.

31. Gracia, M., Theis, S., Proag, A., Gay, G., Benassayag, C. and Suzanne, M. (2019).Mechanical impact of epithelial−mesenchymal transition on epithelial morphogenesis in Drosophila. Nat Commun 10, 2951.

32. Grevengoed, E. E., Loureiro, J. J., Jesse, T. L. and Peifer, M. (2001). Abelson kinase regulates epithelial morphogenesis in Drosophila. J Cell Biol 155, 1185–1198.

33. Grevengoed, E. E., Fox, D. T., Gates, J. and Peifer, M. (2003). Balancing different types of actin polymerization at distinct sites. J Cell Biology 163, 1267–1279.

34. Harmansa, S., Erlich, A., Eloy, C., Zurlo, G. and Lecuit, T. (2022). Growth anisotropy of the extracellular matrix drives mechanics in a developing organ. Biorxiv 2022.07.19.500615.

35. Hayashi, T. and Carthew, R. W. (2004). Surface mechanics mediate pattern formation in the developing retina. Nature 431, 647–652.

36. Hellerman, M. B., Choe, R. H. and Johnson, R. I. (2015). Live-imaging of the Drosophila Pupal Eye. J Vis Exp 52120.

37. Henkemeyer, M. J., Gertler, F. B., Goodman, W. and Hoffmann, F. M. (1987). The Drosophila abelson proto-oncogene homolog: Identification of mutant alleles that have pleiotropic effects late in development. Cell 51, 821–828.

38. Henkemeyer, M., West, S. R., Gertler, F. B. and Hoffmann, F. M. (1990). A novel tyrosine kinase-independent function of Drosophila abl correlates with proper subcellular localization. Cell 63, 949–960.

39. Hilgenfeldt, S., Erisken, S. and Carthew, R. W. (2008). Physical modeling of cell geometric order in an epithelial tissue. Proc National Acad Sci 105, 907–911.

40. Huang, C., Wang, Z., Quinn, D., Suresh, S. and Hsia, K. J. (2018). Differential growth and shape formation in plant organs. Proc National Acad Sci 115, 12359–12364.

41. Irvine, K. D. and Wieschaus, E. (1994). Cell intercalation during Drosophila germband extension and its regulation by pair-rule segmentation genes. Development 120, 827–841.

42. Jodoin, J. N. and Martin, A. C. (2016). Abl suppresses cell extrusion and intercalation during epithelium folding. Mol Biol Cell 27, 2822–2832.

43. Johnson, R. I. (2021). Hexagonal patterning of the Drosophila eye. Dev Biol 478, 173–182.

44. Kafer, J., Hayashi, T., Maree, A. F., Carthew, R. W. and Graner, F. (2007). Cell adhesion and cortex contractility determine cell patterning in the Drosophila retina. Proc Natl Acad Sci U S A 104, 18549–18554.

45. Kannan, R. and Giniger, E. (2017). New perspectives on the roles of Abl tyrosine kinase in axon patterning. Fly 11, 260–270.

46. Kannan, R., Kuzina, I., Wincovitch, S., Nowotarski, S. H. and Giniger, E. (2014). The Abl/Enabled signaling pathway regulates Golgi architecture in Drosophila photoreceptor neurons. Mol Biol Cell 25, 2993–3005.

47. Kannan, R., Song, J.-K., Karpova, T., Clarke, A., Shivalkar, M., Wang, B., Kotlyanskaya, L., Kuzina, I., Gu, Q. and Giniger, E. (2017). The Abl pathway bifurcates to balance Enabled and Rac signaling in axon patterning in Drosophila. Development 144, 487–498.

48. Khalilgharibi, N., Fouchard, J., Asadipour, N., Barrientos, R., Duda, M., Bonfanti, A., Yonis, A., Harris, A., Mosaffa, P., Fujita, Y., et al. (2019). Stress relaxation in epithelial monolayers is controlled by the actomyosin cortex. Nat Phys 15, 839–847.

49. Kiehart, D. P., Galbraith, C. G., Edwards, K. A., Rickoll, W. L. and Montague, R. A. (2000). Multiple Forces Contribute to Cell Sheet Morphogenesis for Dorsal Closure in Drosophila. J Cell Biology 149, 471–490.

50. Lee, Y. S. and Carthew, R. W. (2003). Making a better RNAi vector for Drosophila: use of intron spacers. Methods 30, 322–329.

51. Liebl, E. C., Forsthoefel, D. J., Franco, L. S., Sample, S. H., Hess, J. E., Cowger, J. A., Chandler, M. P., Shupert, A. M. and Seeger, M. A. (2000). Dosage-sensitive, reciprocal genetic interactions between the Abl tyrosine kinase and the putative GEF trio reveal trio’s role in axon pathfinding. Neuron 26, 107–118.

52. Lin, T. Y., Huang, C. H., Kao, H. H., Liou, G. G., Yeh, S. R., Cheng, C. M., Chen, M. H., Pan, R. L. and Juang, J. L. (2009). Abi plays an opposing role to Abl in Drosophila axonogenesis and synaptogenesis. Development 136, 3099–3107.

53. Longley, R. L. and Ready, D. F. (1995). Integrins and the Development of Three-Dimensional Structure in the Drosophila Compound Eye. Dev Biol 171, 415–433.

54. Mao, Y. and Baum, B. (2015). Tug of war—The influence of opposing physical forces on epithelial cell morphology. Dev Biol 401, 92–102.

55. Mao, Y., Tournier, A. L., Hoppe, A., Kester, L., Thompson, B. J. and Tapon, N. (2013). Differential proliferation rates generate patterns of mechanical tension that orient tissue growth. Embo J 32, 2790–2803.

56. Martin, A. C. and Goldstein, B. (2014). Apical constriction: themes and variations on a cellular mechanism driving morphogenesis. Development 141, 1987–1998.

57. Martin, A. C., Kaschube, M. and Wieschaus, E. F. (2009). Pulsed contractions of an actin–myosin network drive apical constriction. Nature 457, 495–499.

58. Matsuo, T., Takahashi, K., Suzuki, E. and Yamamoto, D. (1999). The Canoe protein is necessary in adherens junctions for development of ommatidial architecture in the Drosophila compound eye. Cell Tissue Res 298, 397–404.

59. Mishra, A. K., Tsachaki, M., Rister, J., Ng, J., Celik, A. and Sprecher, S. G. (2013). Binary Cell Fate Decisions and Fate Transformation in the Drosophila Larval Eye. Plos Genet 9, e1004027.

60. Nagarkar-Jaiswal, S., Lee, P.-T., Campbell, M. E., Chen, K., Anguiano-Zarate, S., Gutierrez, M. C., Busby, T., Lin, W.-W., He, Y., Schulze, K. L., et al. (2015). A library of MiMICs allows tagging of genes and reversible, spatial and temporal knockdown of proteins in Drosophila. Elife 4, e05338.

61. O’Donnell, M. P. and Bashaw, G. J. (2013). Distinct functional domains of the Abelson tyrosine kinase control axon guidance responses to Netrin and Slit to regulate the assembly of neural circuits. Development 140, 2724–2733.

62. Paré, A. C. and Zallen, J. A. (2020). Cellular, molecular, and biophysical control of epithelial cell intercalation. Curr Top Dev Biol 136, 167–193.

63. Pellikka, M., Tanentzapf, G., Pinto, M., Smith, C., McGlade, C. J., Ready, D. F. and Tepass, U. (2002). Crumbs, the Drosophila homologue of human CRB1/RP12, is essential for photoreceptor morphogenesis. Nature 416, 143–9.

64. Perez-Vale, K. Z. and Peifer, M. (2020). Orchestrating morphogenesis: building the body plan by cell shape changes and movements. Development 147, dev191049.

65. Pham, H., Yu, H. and Laski, F. A. (2008). Cofilin/ADF is required for retinal elongation and morphogenesis of the Drosophila rhabdomere. Dev Biol 318, 82–91.

66. Pickup, A. T., Lamka, M. L., Sun, Q., Yip, M. L. R. and Lipshitz, H. D. (2002). Control of photoreceptor cell morphology, planar polarity and epithelial integrity during Drosophila eye development. Development 129, 2247–2258.

67. Popkova, A., Rauzi, M. and Wang, X. (2021). Cellular and Supracellular Planar Polarity: A Multiscale Cue to Elongate the Drosophila Egg Chamber. Frontiers Cell Dev Biology 9, 645235.

68. Raghu, P., Coessens, E., Manifava, M., Georgiev, P., Pettitt, T., Wood, E., Garcia-Murillas, I., Okkenhaug, H., Trivedi, D., Zhang, Q., et al. (2009). Rhabdomere biogenesis in Drosophila photoreceptors is acutely sensitive to phosphatidic acid levels. J Cell Biol 185, 129–145.

69. Ready, D. F. (2002). Drosophila compound eye morphogenesis: Blind mechanical engineers? K. Moses *(Ed.)* Results and Problems in Cell Differentiation, Springer*-* Verlag.

70. Ready, D. F. and Chang, H. C. (2021). Calcium waves facilitate and coordinate the contraction of endfeet actin stress fibers in Drosophila interommatidial cells. Development 148,.

71. Ready, D. F., Hanson, T. E. and Benzer, S. (1976a). Development of the Drosophila retina, a neurocrystalline lattice. Dev Biol 53, 217–240.

72. Ready, D. F., Hanson, T. E. and Benzer, S. (1976b). Development of the Drosophila retina, a neurocrystalline lattice. Dev Biol 53, 217–240.

73. Rebocho, A. B., Southam, P., Kennaway, J. R., Bangham, J. A. and Coen, E. (2017). Generation of shape complexity through tissue conflict resolution. Elife 6, e20156.

74. Roellig, D., Theis, S., Proag, A., Allio, G., Bénazéraf, B., Gros, J. and Suzanne, M. (2022). Force-generating apoptotic cells orchestrate avian neural tube bending. Dev Cell.

75. Rogers, E. M., Allred, S. C. and Peifer, M. (2021). Abelson kinase’s intrinsically disordered region plays essential roles in protein function and protein stability. Cell Commun Signal 19, 27.

76. Sherrard, K., Robin, F., Lemaire, P. and Munro, E. (2010). Sequential Activation of Apical and Basolateral Contractility Drives Ascidian Endoderm Invagination. Curr Biol 20, 1499–1510.

77. Shindo, A. (2018). Models of convergent extension during morphogenesis. Wiley Interdiscip Rev Dev Biology 7, e293.

78. Shyer, A. E., Tallinen, T., Nerurkar, N. L., Wei, Z., Gil, E. S., Kaplan, D. L., Tabin, C. J. and Mahadevan, L. (2013). Villification: How the Gut Gets Its Villi. Science 342, 212–218.

79. Signore, S. J. D., Cilla, R. and Hatini, V. (2018). The WAVE Regulatory Complex and Branched F-Actin Counterbalance Contractile Force to Control Cell Shape and Packing in the Drosophila Eye. Dev Cell 44, 471–483.e4.

80. Silva, S. M. da and Vincent, J.-P. (2007). Oriented cell divisions in the extending germband of Drosophila. Development 134, 3049–3054.

81. Singh, J., Yanfeng, W. A., Grumolato, L., Aaronson, S. A. and Mlodzik, M. (2010). Abelson family kinases regulate Frizzled planar cell polarity signaling via Dsh phosphorylation. Genes Dev 24, 2157–2168.

82. Stooke-Vaughan, G. A. and Campàs, O. (2018). Physical control of tissue morphogenesis across scales. Curr Opin Genet Dev 51, 111–119.

83. Suarez, C. and Kovar, D. R. (2016). Internetwork competition for monomers governs actin cytoskeleton organization. Nat Rev Mol Cell Bio 17, 799–810.

84. Sui, L., Alt, S., Weigert, M., Dye, N., Eaton, S., Jug, F., Myers, E. W., Jülicher, F., Salbreux, G. and Dahmann, C. (2018). Differential lateral and basal tension drive folding of Drosophila wing discs through two distinct mechanisms. Nat Commun 9, 4620.

85. Tamada, M., Farrell, D. L. and Zallen, J. A. (2012). Abl regulates planar polarized junctional dynamics through beta-catenin tyrosine phosphorylation. Dev Cell 22, 309–319.

86. Warga, R. M. and Kimmel, C. B. (1990). Cell movements during epiboly and gastrulation in zebrafish. Development 108, 569–580.

87. Wills, Z., Bateman, J., Korey, C. A., Comer, A. and Vactor, D. V. (1999). The Tyrosine Kinase Abl and Its Substrate Enabled Collaborate with the Receptor Phosphatase Dlar to Control Motor Axon Guidance. Neuron 22, 301–312.

88. Wilson, P. and Keller, R. (1991). Cell rearrangement during gastrulation of Xenopus: direct observation of cultured explants. Development 112, 289–300.

89. Wolff, T. and Ready, D. F. (1993). Pattern Formation in the Drosophila Retina. Cold Spring Harbor Laboratory Press 1277–1325.

90. Wyatt, T. P. J., Fouchard, J., Lisica, A., Khalilgharibi, N., Baum, B., Recho, P., Kabla, A. J. and Charras, G. T. (2020). Actomyosin controls planarity and folding of epithelia in response to compression. Nat Mater 19, 109–117.

91. Xiong, W. and Rebay, I. (2011). Abelson tyrosine kinase is required for Drosophila photoreceptor morphogenesis and retinal epithelial patterning. Dev Dyn 240, 1745–1755.

92. Xiong, W., Dabbouseh, N. M. and Rebay, I. (2009). Interactions with the abelson tyrosine kinase reveal compartmentalization of eyes absent function between nucleus and cytoplasm. Dev Cell 16, 271–279.

93. Xiong, W., Morillo, S. A. and Rebay, I. (2013). The Abelson tyrosine kinase regulates Notch endocytosis and signaling to maintain neuronal cell fate in Drosophila photoreceptors. Development 140, 176–184.

94. Yang, Q., Roiz, D., Mereu, L., Daube, M. and Hajnal, A. (2017). The Invading Anchor Cell Induces Lateral Membrane Constriction during Vulval Lumen Morphogenesis in C. elegans. Dev Cell 42, 271–285.e3.

95. Yevick, H. G., Miller, P. W., Dunkel, J. and Martin, A. C. (2019). Structural Redundancy in Supracellular Actomyosin Networks Enables Robust Tissue Folding. Dev Cell 50, 586–598.e3.

96. Yu, H. H. and Zallen, J. A. (2020). Abl and Canoe/Afadin mediate mechanotransduction at tricellular junctions. Science 370, eaba5528.

97. Zallen, J. A. and Wieschaus, E. (2004). Patterned Gene Expression Directs Bipolar Planar Polarity in Drosophila. Dev Cell 6, 343–355.

